# CTCF-RNA interactions orchestrate cell-specific chromatin loop organization

**DOI:** 10.1101/2025.03.19.643339

**Authors:** Kimberly Lucero, Sungwook Han, Pin-Yao Huang, Xiang Qiu, Esteban O. Mazzoni, Danny Reinberg

**Affiliations:** Department of Cell Biology and Regenerative Medicine, New York University Langone Medical Center, New York, NY, USA; Department of Biochemistry and Molecular Pharmacology, New York University Langone Medical Center, New York, NY, USA; Sylvester Comprehensive Cancer Center, Department of Human Genetics, University of Miami Miller School of Medicine, Miami, FL, USA; Howard Hughes Medical Institute, University of Miami Miller School of Medicine, Miami, FL, USA

**Keywords:** CTCF, chromatin loops, genome organization, RNA binding, gene expression, embryonic stem cells, neural progenitor cells

## Abstract

CCCTC-binding factor (CTCF) is essential for chromatin organization. CTCF interacts with endogenous RNAs, and deletion of its ZF1 RNA-binding region (ΔZF1) disrupts chromatin loops in mouse embryonic stem cells (ESCs). However, the functional significance of CTCF-ZF1 RNA interactions during cell differentiation is unknown. Using an ESC-to-neural progenitor cell (NPC) differentiation model, we show that CTCF-ZF1 is crucial for maintaining cell-type-specific chromatin loops. Expression of CTCF-ΔZF1 leads to disrupted loops and dysregulation of genes within these loops, particularly those involved in neuronal development and function. We identified NPC-specific, CTCF-ZF1 interacting RNAs. Truncation of two such coding RNAs, *Podxl* and *Grb10*, disrupted chromatin loops *in cis*, similar to the disruption seen in CTCF-ΔZF1 expressing NPCs. These findings underscore the inherent importance of CTCF-ZF1 RNA interactions in preserving cell-specific genome structure and cellular identity.

**HIGHLIGHTS:**

- CTCF loop anchors induced after differentiation are disrupted in the ΔZF1 RNA-binding mutant.
- Loop loss in the ΔZF1 mutant is independent of its DNA binding and protein interactions.
- Chromatin loop loss is associated with gene dysregulation.
- Truncation of cell-specific, CTCF-ZF1-interacting RNAs disrupts chromatin loops *in cis*.

**GRAPHICAL ABSTRACT:** 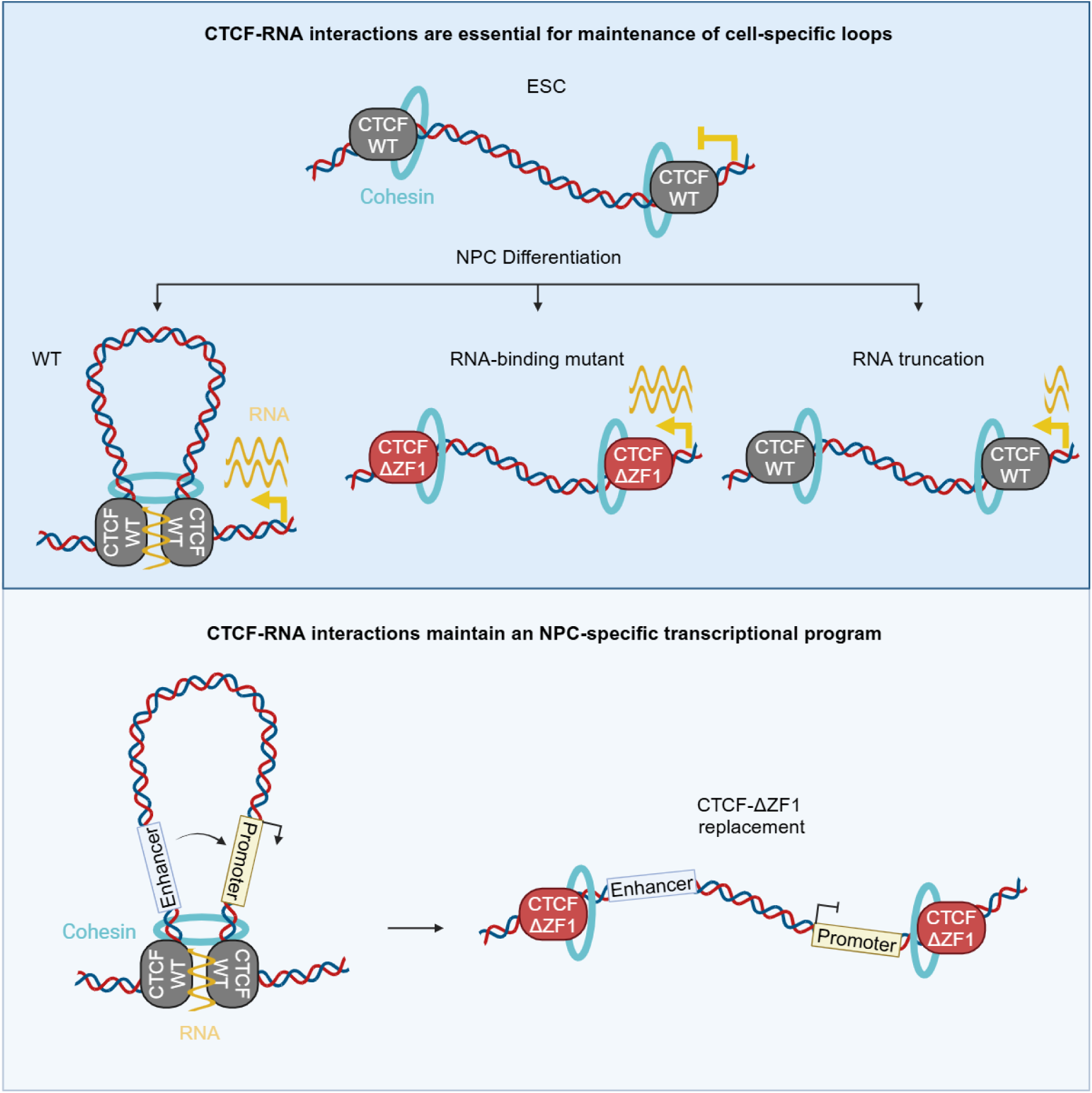

## INTRODUCTION

One of the long-standing questions in biology is how a single cell can give rise to multiple cell types. Despite having the same genome, gene expression varies from one cell type to another, resulting in the diverse tissues that comprise a multicellular organism. The non-random structural organization of the genome is crucial for maintaining cell-type-specific transcriptional programs that establish identity and proper function^1–3^. Indeed, disruption of chromatin structure can lead to oncogenesis or developmental abnormalities^4–7^.

The structure of the genome is organized hierarchically. Interphase chromosomes occupy well-defined territories within the nuclear space^8^. Within these chromosome territories are A and B compartments, corresponding to euchromatin and heterochromatin, respectively^9^. Within compartments are fundamental structures, called Topologically Associating Domains (TADs), which are defined as contiguous chromatin that is more frequently interacting with itself compared to chromatin outside the TAD^10,11^. TADs are insulated regions of chromatin, which are composed of either actively transcribed regions, or silent chromatin domains^12^. And finally, colocalized at or within TAD boundaries are chromatin loops, defined as genomic regions that are in closer physical proximity to each other than to their intervening sequences^13^. TAD boundaries that are demarcated by chromatin loops occur frequently and are referred to as ‘loop domains’^13^.

Central to the structural maintenance of chromatin loops (as well as TADs) in the mammalian system is CCCTC-binding factor (CTCF)^14^, which is ubiquitously expressed and binds with high affinity to a conserved DNA motif^15,16^. CTCF binding sites are enriched at most chromatin loop anchors^13^, and are important for maintaining chromatin loops^17^. Chromatin loops bound by CTCF at their base (‘CTCF loop anchors’) are frequently found in close proximity to genomic regions enriched for active histone modifications, RNA polymerase II, and transcription start sites (TSS)^18^. These anchors play multi-faceted and context-specific roles in transcriptional regulation by: 1) bringing promoters close to distal enhancers to promote transcription^19^, 2) preventing ectopic contacts between promoters and enhancers not within the same loop domain^5,12^, and 3) acting as insulators or barriers to the spreading of repressive histone modifications^6,7^.

The formation of CTCF loop anchors involves CTCF interaction with the cohesin complex, composed of Rad21, Smc3, Smc1a, and Stag1/Stag2^20,21^. Chromatin loops form apparently through a loop extrusion mechanism, whereby cohesin tethers two regions together and extrudes chromatin until it contacts two distant CTCF-bound sites in convergent orientation^22–28^. Cohesin accessory proteins, such as Nipbl-Mau2, Wapl, Pds5a, and Pds5b, regulate cohesin loading, unloading, and processivity, thus modulating chromatin loop formation^29–31^.

While the loop extrusion model is an elegant and widely accepted explanation for how chromatin loops are formed, some observations regarding chromatin loops remain unexplained. Although most chromatin loops are bound by CTCF, there are more CTCF binding sites than loop anchors, and most CTCF binding sites are not at chromatin loops^10,21^. It remains unclear why certain CTCF binding sites function as loop anchors while others function in transcription. Additionally, CTCF loop anchors are dynamic and can change upon differentiation, such as during the differentiation of mouse embryonic stem cells (ESCs) into neural progenitor cells (NPCs)^32,33^. The sites to which CTCF actively binds are largely invariant between cell types^34^, yet they exhibit differential chromatin interactions^32,35^. Thus, the reorganization of CTCF loop anchors upon differentiation is not well understood and likely involves cell-specific regulatory elements that cooperate with CTCF. Indeed, it was previously found that, in addition to CTCF, other proteins such as Maz, Patz1, and Znf263 modulate insulation and gene expression in a cell-specific manner^36–38^.

In this study, we examined the role of cell-specific CTCF-RNA interactions in chromatin loop formation based on the following evidence. CTCF comprises of a central zinc-finger domain with 11 zinc fingers (ZF). The central ZF3-7 directly interact with the 15-bp core DNA motif and are crucial for CTCF DNA-binding^16,39^. The peripheral ZF1, ZF10, and a C-terminal region (RBRi) are RNA-binding domains^40–43^. CTCF interacts with thousands of endogenous RNAs^44^. Notably, a deletion in ZF1 of CTCF (ΔZF1) thwarts the integrity of some chromatin loops *in vivo*. Yet, importantly, deletion or point mutation in ZF1 do not affect CTCF DNA-binding activity *in vitro*^45^, and give rise to a modest loss *in vivo*^39^. A deletion in ZF10 of CTCF (ΔZF10) also results in decreased integrity of chromatin loops, albeit to a much lesser degree compared to ΔZF1^41^. ΔZF1 and ΔZF10 mutants display defects in self-association compared to WT^41^, which may explain the observed loss of chromatin loops. Similarly, a deletion in RBRi (ΔRBRi) leads to loss of chromatin loops, as well as decreased CTCF protein self-clustering *in vitro*^40^ and *in vivo*^43^. In addition to its role in facilitating CTCF-CTCF interactions, RNA has been shown to either facilitate or decrease CTCF binding to chromatin and thereby influence chromatin architecture and transcription^44–50^.

Given the evidence above, we focused on elucidating the broader role and importance of CTCF-ZF1 in chromatin loop maintenance. Using an *in vitro* model system of ESC to NPC differentiation^51^, we sought to answer the following questions: (1) are CTCF-ZF1 RNA-interactions required for the maintenance of cell-type-specific chromatin loops?, (2) how does the ΔZF1 mutant affect gene expression within these loops?, and (3) are there cell-type-specific CTCF-ZF1 RNA-interactions, and if so, are they important in maintaining cell-type-specific chromatin loops? Using a CTCF-AID2 degron system^52^ coupled with rescue experiments involving doxycycline (dox)-inducible CTCF, either WT or ΔZF1, we uncovered a vital role of CTCF-RNA interactions in chromatin loop organization. Our findings highlight that CTCF-ZF1 interactions with RNA are central to the integrity of cell-specific genome structure and cellular identity.

## RESULTS

### Acute CTCF depletion and rescue using the AID2 system

To determine the role of CTCF-ZF1 in genome organization, we generated mouse embryonic stem cell lines (ES-E14TG2a) with an endogenous, homozygous deletion of CTCF-ZF1 (amino acids 264-277; ΔZF1) (**Fig. S1A, S1B**). The ΔZF1 ESCs had similar CTCF and Rad21 protein levels as WT, but exhibited decreased protein levels of the pluripotency gene Oct3/4 (**Fig. S1C**), suggesting that the mutant would have differentiation defects. We differentiated the ΔZF1 ESCs into NPCs following an established protocol^51^, and assessed differentiation efficiency by immunostaining with Sox1, an NPC-specific marker^51^. The ΔZF1 mutant exhibited inefficient differentiation, with a reduced percentage (**Fig. S1D**) and total count of Sox1+ cells (**Fig. S1E**), indicating that the CTCF-ZF1 region is crucial for cellular differentiation.

Given the differentiation challenges with ESCs expressing ΔZF1 constitutively, we instead developed ESC lines having an AID2 degron system for CTCF, coupled with dox-inducible CTCF rescues (**Fig. 1A, S2A**). This system allows for the degradation of endogenous CTCF and the inducible expression of tagged rescue CTCF variants, either wild-type (WT) or ΔZF1, in a temporally controlled manner post-differentiation into NPCs.

**Figure 1.**
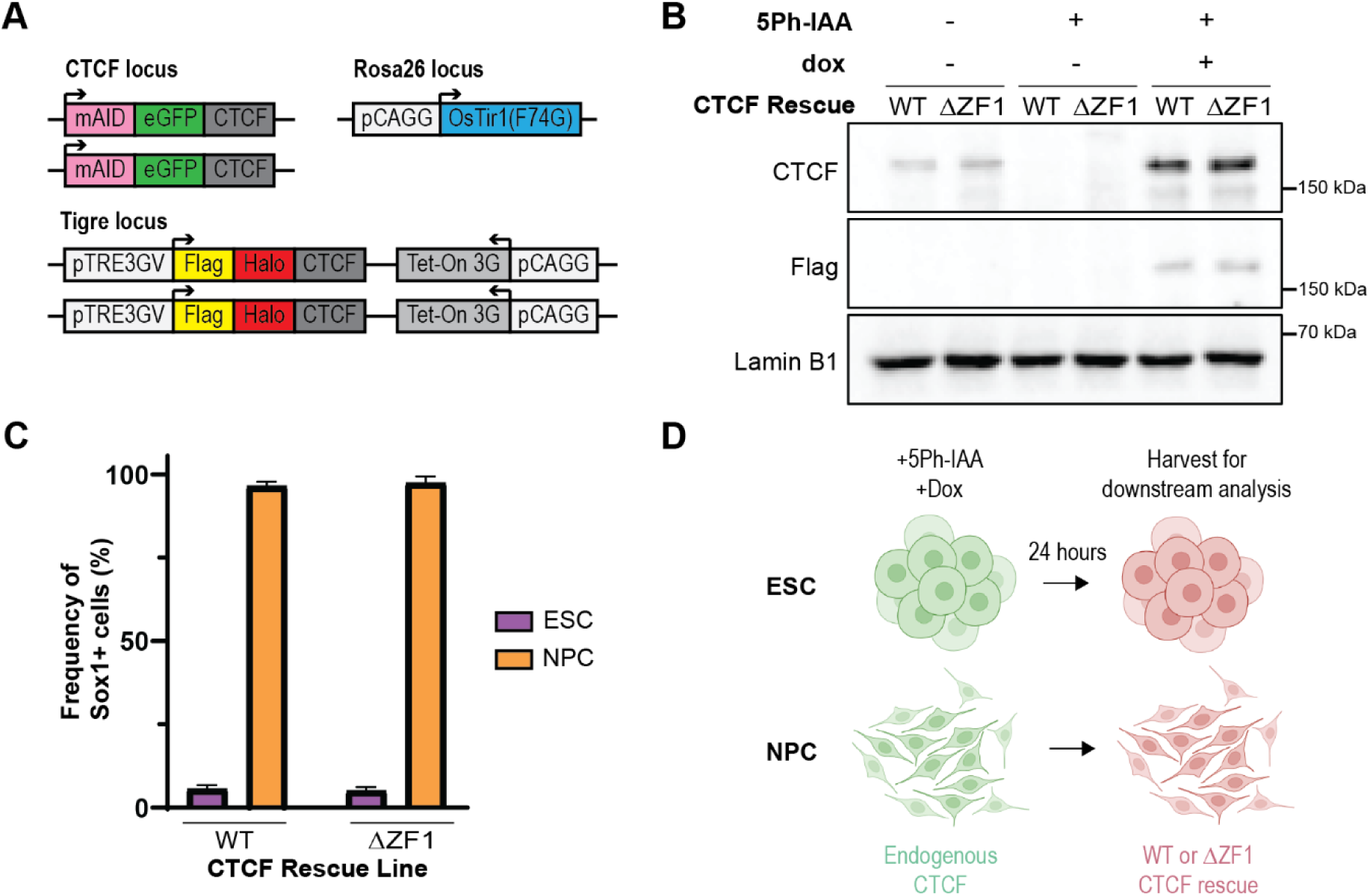
Acute CTCF depletion and rescue using the AID2 system. (A) Schematic showing CTCF-AID2 degron and dox-inducible rescue system generated in ESCs. (B) Western blot of CTCF-AID2 and rescue lines after 24 hours with no treatment, or treatment with 5Ph-IAA, or 5Ph-IAA and dox. (C) CTCF-AID2 and dox-inducible rescue lines were differentiated into NPCs for 2 days. Cells were immunostained with anti-Sox1 antibody and percentages of Sox1+ cells were quantified by flow cytometry analysis. Neither 5-Ph-IAA nor dox was added. Data are represented as mean ± SEM, N=2. (D) Diagram of experimental approach. ESCs or Day 2 NPCs were treated with 5-Ph-IAA and dox, then harvested for downstream analysis after 24 hours. Image generated using Biorender (biorender.com).

To generate the CTCF-AID2 degron in the ES-E14TG2a background, we used CRISPR-Cas9-based gene editing^53^ to tag both endogenous CTCF alleles with mAID-eGFP at the N-terminus. We then inserted OsTir1(F74G) into the Rosa26 locus to facilitate inducible degradation. OsTir1(F74G) interacts with mAID in the presence of the auxin analog 5-Ph-IAA, leading to ubiquitination by E3 ligase and subsequent proteasome-dependent degradation of mAID-eGFP-tagged CTCF^52^. We then integrated dox-inducible Flag-Halo-tagged versions of CTCF, either WT (control) *or* ΔZF1 into the Tigre locus of the CTCF-AID2 degron line.

First, we assessed the timing of CTCF degradation in ESCs using eGFP fluorescence (mAID-eGFP-CTCF). As expected, 100% of CTCF-AID2 cells strongly expressed eGFP, and the addition of 5Ph-IAA led to a near-complete loss of eGFP fluorescence within 1 hour (**Fig. S2B**). Next, we tested the efficiency of rescue expression by conjugating the Halo portion of the tagged CTCF to a JF646-fluorescent HaloTag. Almost 100% of cells expressed CTCF, either WT or ΔZF1, at 24 hours after dox treatment (**Fig. S2C**). Western blot analysis also confirmed degradation of mAID-eGFP-CTCF, and showed similar Flag-Halo-CTCF protein levels between the WT and ΔZF1 rescues (**Fig. 1B**). We verified that the WT and ΔZF1 rescue lines expressing tagged endogenous CTCF (without 5Ph-IAA or dox treatment) showed Sox1 expression in over 90% of cells by Day 2 of NPC differentiation (**Fig. 1C**). For the experimental set-up, we maintained ESCs in culture, while differentiating the same lines into NPCs in parallel. ESCs and NPCs were then treated with 5Ph-IAA and dox and harvested after 24 hours for analysis (**Fig. 1D**). We limited the rescue expression to 24 hours to capture more direct structural effects and minimize indirect effects from cascading gene expression changes due to mutant expression. Using this CTCF-AID2 and dox-inducible rescue system, we next investigated the role of CTCF-ZF1 RNA interactions in both differentiated NPCs and their parental ESCs.

### CTCF loop anchors induced after differentiation are disrupted in CTCF-ΔZF1

To determine the role of CTCF-ZF1 on genome organization, we first assessed the impact of the ΔZF1 mutation on CTCF loop anchors in ESCs and NPCs expressing the ΔZF1 mutant rescue (ESC-ΔZF1 and NPC-ΔZF1, respectively). We performed Micro-C^54,55^ and used FitHiC2^56^ to identify chromatin loops at a 5-kb resolution (FDR ≤ 0.01). To specifically identify CTCF loop anchors, we performed CTCF ChIP-seq to determine CTCF binding sites and overlapped called ChIP-seq peaks with chromatin loop boundaries. We observed that ∼40% of all called loops had at least one occupied CTCF binding site at one of the anchors and our subsequent analysis focused on this subset. We performed an aggregate peak analysis (APA), which quantifies the aggregate interaction signal of multiple chromatin loops.^13^ We plotted the APA of CTCF WT loop anchors in WT and compared it to the APA of ΔZF1. Consistent with previous findings^41^, we observed a decrease in CTCF loop anchors in ESC-ΔZF1 (**Fig. 2A**). A similar reduction was also observed in NPC-ΔZF1 (**Fig. 2B**). We then categorized the WT loops based on whether they were significantly decreased (log_2_ fold-change ≤-1) or not in the mutant, labeling them as ΔZF1-lost and ΔZF1-retained, respectively. We found that 40% (1,504/3,860) of CTCF loop anchors in ESCs and 50% (2,238/4,495) in NPCs were lost in the ΔZF1 mutant (**Fig. 2C**).

**Figure 2.**
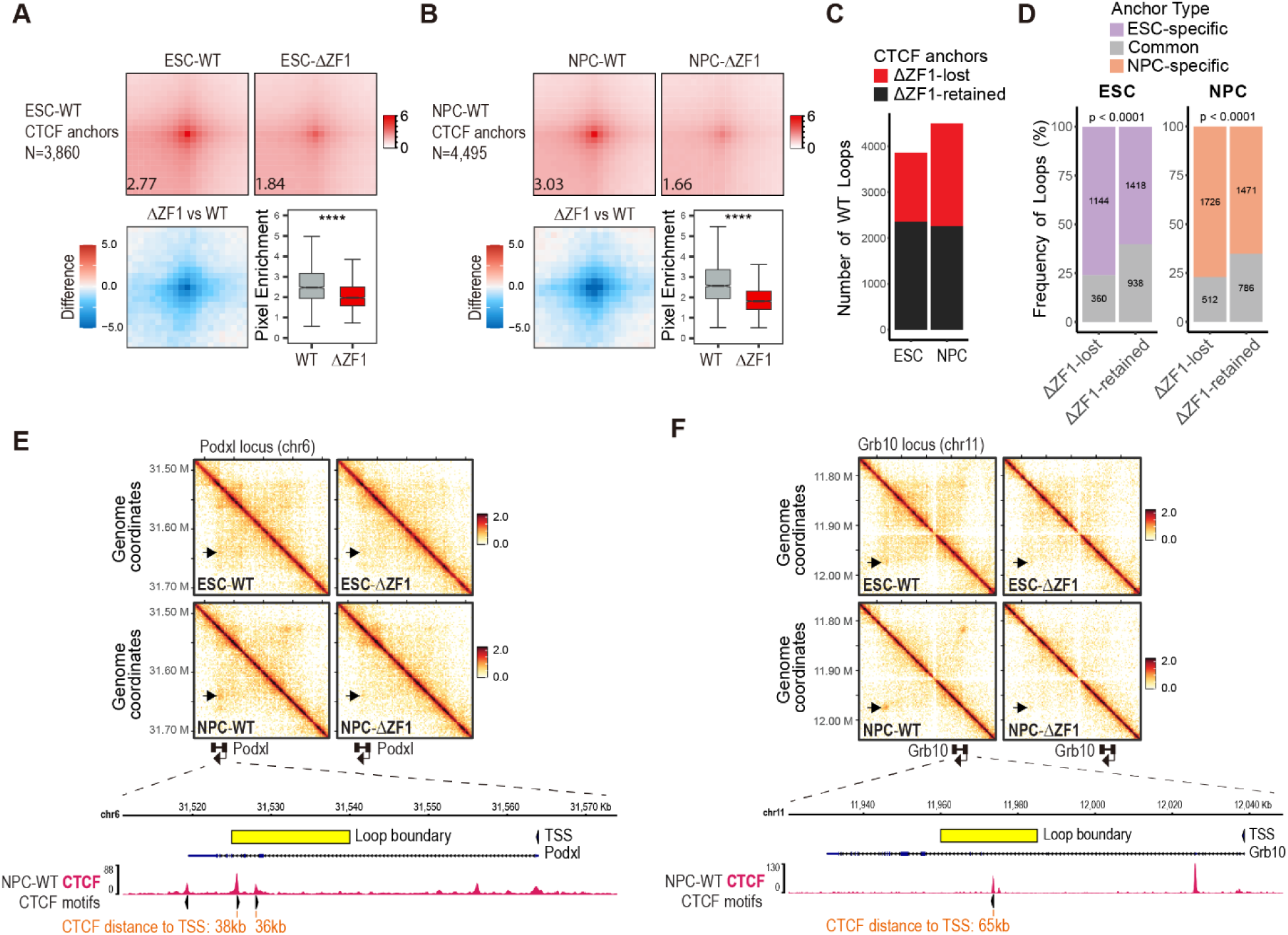
CTCF loop anchors induced after differentiation are disrupted in the ΔZF1 mutant. (A, B) Aggregate peak analysis (APA) plots (top) of all WT CTCF loop anchors in (A) ESCs and (B) NPCs, comparing WT and ΔZF1. The numbers indicate the APA score. Lower-left is the difference in APA enrichment between ΔZF1 and WT. Lower-right are box plots quantifying the pixel enrichment of each loop, comparing WT and ΔZF1. p-values were determined using Wilcoxon signed-rank test. **** p-value < 1×10^-100^. (C) Bar plots showing the number of WT CTCF loop anchors and the proportions that are lost or retained in the ΔZF1 mutant. Loops were classified as ΔZF1-lost if they were found to be significant only in WT but not in ΔZF1 (q-value cutoff of 0.01), and if contact counts had log_2_ fold-change ≤-1 (see Methods). (D) Bar plot showing the proportions of cell-type-specific and common loops among ΔZF1-lost or ΔZF1-retained loops. The numbers indicate the absolute numbers of loops. p-values were determined using Fisher’s Exact test. (E, F) Micro-C contact heatmaps (top) at the (E) *Podxl* and (F) *Grb10* loci. Arrows point to NPC-specific CTCF-bound loops. Below the heatmaps is the view of one of the loop boundaries (indicated by the yellow bar), which is located in the gene body of (E) *Podxl* and (F) *Grb10*. Below the gene annotation are CTCF ChIP-seq tracks, and the annotation of conserved CTCF motifs with their orientation. The distance of the CTCF loop boundary sites to the gene TSS is indicated.

We then evaluated the impact of ΔZF1 on higher-order chromatin organization. We used principal component analysis (PCA)^9^ to identify the spatial segregation of A and B compartments, corresponding to positive and negative values of the first eigenvector, respectively. Our analysis revealed that the compartments in ESC-ΔZF1 and NPC-ΔZF1 were largely similar to those in WT cells, with only ∼3% of compartments showing a shift from A-to-B or B-to-A (**Fig. S3A**). We found a high correlation between the first eigenvectors of WT and ΔZF1 (R²=0.98) (**Fig. S3B, S3C**), and the absence of noticeable changes on the plaid pattern observed in Micro-C heatmaps (**Fig. S3D, S3E**). These results are consistent with previous studies showing that compartmentalization is CTCF-independent^17,21^.

Chromatin loops are frequently colocalized at TAD boundaries^13^, and the loop extrusion model proposes that TADs emerge from multiple dynamically formed loops during extrusion^24,26^. Given the widespread reduction in loops, we investigated whether TADs were also affected in ΔZF1. We used Arrowhead^57^ to identify TADs and performed Aggregate TAD Analysis (ATA)^58^ to average the results of all TADs. Comparison of ATA between WT and ΔZF1 revealed a decrease in long-range interactions within TADs in ΔZF1 (**Fig. S4A, S4B**). The decrease in interactions was most prominently observed at TAD boundaries (**Fig. S4A, S4B**). To compare the level of interaction separation at TAD boundaries, we calculated diamond insulation scores^59^, which quantify the degree of interactions with neighboring regions. A lower insulation score indicates a stronger TAD boundary. We plotted the insulation score profiles for WT and ΔZF1 centered on CTCF ChIP-seq peaks and observed an increase in insulation scores in ΔZF1, indicating weakened TAD boundaries (**Fig. S4C, S4D**). These results show that the CTCF-ZF1 RNA-binding region is crucial for maintaining not just CTCF loop anchors, but also TADs.

To ascertain the integrity of cell-specific loops in the presence of the ΔZF1 mutant, we first identified cell-specific and ESC/NPC common (or constitutive) loops. As expected, the differentiation of WT ESCs into WT NPCs resulted in differential CTCF loop anchors (**Fig. S5A**). Notably, the presence of CTCF and cohesin at DNA-binding sites were largely cell-type-invariant (**Fig. S5B, S5C, S5D**), and binding at cell-specific loop anchors was similar between the cell types (**Fig. S5E, S5F, S5G**). We then investigated whether cell-specific loop subsets were affected in ΔZF1. We found in ESCs that cell-specific CTCF anchors constituted 76% (1,144/1,504) of ΔZF1-lost anchors and 60% (1,418/2,356) of ΔZF1-retained anchors (**Fig. 2D**). Similarly, in NPCs, cell-specific CTCF anchors comprised 77% (1,726/2,238) of ΔZF1-lost anchors and 65% (1,417/2,257) of ΔZF1-retained anchors (**Fig. 2D**). Statistical analysis (Fisher’s exact test) indicated a significant difference in the distribution of cell-specific loops. The likelihood (odds ratio) of losing a cell-specific loop compared to losing a common loop was 2.1 times higher in ESCs, and 1.8 times higher in NPCs. We show examples of NPC-specific, ΔZF1-lost loops at the *Podxl* (**Fig. 2E**) and *Grb10* (**Fig. 2F**) loci. These loci are illustrated as our subsequent studies will focus on the function of CTCF interaction with *Podxl* and *Grb10* RNAs. We highlight one of the loop boundaries at each locus, showing that the CTCF loop anchor coordinates are located within the *Podxl* (**Fig. 2E**) and *Grb10* (**Fig. 2F**) gene body.

Based on these results, we concluded that the CTCF-ZF1 RNA-binding region plays an essential role in maintaining loops not only in ESCs, but also in the case of the re-organized CTCF loop anchors that arise after differentiation into NPCs.

### ΔZF1 loop loss is not due to loss of CTCF-DNA or CTCF-cohesin interactions

ΔZF1 has been shown to have an RNA-binding defect^41^, yet the loss of CTCF anchors may be due to other functional deficits arising from the deletion of ZF1. Therefore, we first probed whether the loss of CTCF anchors might arise from a decrease in CTCF DNA-binding activity and/or changes in CTCF-protein interactions, particularly that of CTCF-cohesin.

To address DNA binding activity, we performed CTCF ChIP-seq on WT and ΔZF1 rescues. ChIP-seq peaks were called using MACS2^60^, and differential peaks were identified using DiffBind^61^. We observed minimal changes in CTCF ChIP-seq peaks in the ΔZF1 mutant, with only 0.02% (8/38,106) of peaks significantly decreased in ESCs (**Fig. 3A**) and 0.18% (58/31,354) significantly decreased in NPCs (**Fig. 3B**). Importantly, we then quantified the proportion of ΔZF1-lost loops that had decreased CTCF DNA-binding at the anchors. We found that none of the 1,504 ESC-ΔZF1-lost and none of the 2,356 ESC-ΔZF1-retained anchors had changes in CTCF binding. In accordance, only 9 of 2,238 NPC-ΔZF1-lost anchors, and 6 of 2,257 NPC-ΔZF1-retained anchors, showed decreased CTCF binding (**Fig. 3C**). Furthermore, ChIP-seq heatmaps showing the enrichment of CTCF at the anchors of ΔZF1-lost and ΔZF1-retained loops revealed no gross changes in ΔZF1 (**Fig. 3D, 3E**).

**Figure 3.**
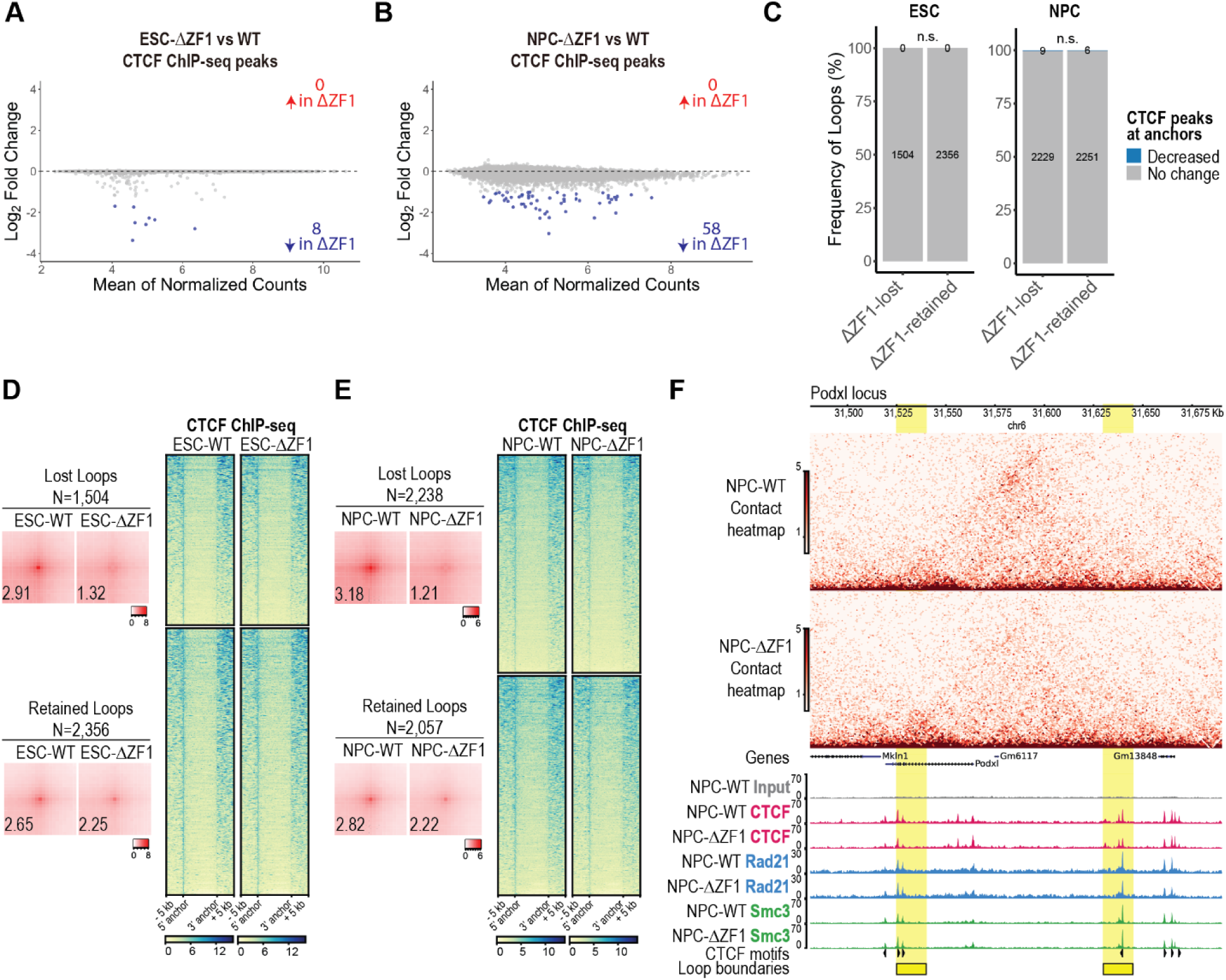
ΔZF1 loop loss is not due to changes in CTCF chromatin binding. (A, B) DiffBind MA plot of differentially called peaks comparing ΔZF1 vs WT in (A) ESCs and (B) NPCs. Adjusted p-value cutoff: ≤ 0.05, log_2_ fold-change cutoff: ≥ 1, ≤-1, N=4. (C) Bar plot showing the proportion of loops that exhibit decreased or no change in CTCF chromatin binding at the anchors. Numbers indicate the absolute numbers of loops. p-values were determined using Fisher’s Exact test (n.s. = not significant). (D, E) (Left) The APA plots in (D) ESCs and (E) NPCs show the comparison between WT and ΔZF1 for the lost and retained loop subsets. Numbers indicate APA scores. (Right) CTCF ChIP-seq heatmaps comparing CTCF chromatin binding in (D) ESC-WT and ESC-ΔZF1 and (E) NPC-WT and NPC-ΔZF1. Each row is a loop anchor coordinate, and the heatmap is clustered based on whether the anchor is ΔZF1-lost or ΔZF1-retained. (F) Micro-C contact heatmap (above) and ChIP-seq tracks for CTCF and cohesin subunits, Rad21 and Smc3 (below) at a representative *Podxl* locus. Loop boundaries are highlighted in yellow. CTCF motifs with their orientation are annotated.

We next investigated whether ΔZF1 could lead to a loss in CTCF-protein interactions. We performed native anti-Flag immunoprecipitation of Flag-Halo-tagged rescue CTCF from the chromatin fraction, followed by mass spectrometry (ChIP-MS) on WT and ΔZF1 rescues. As expected, we detected all the cohesin subunits as CTCF-protein interactors (**Fig. S6A, Table S1**). Moreover, we did not observe any differential protein interactions between ΔZF1 and WT (**Fig. S6A, Table S1**). This finding contrasts with that of another CTCF-ZF mutant (ΔZF10), with which a handful of differential interactions had been detected (**Fig. S6B, Table S2**). Consequently, we did not further investigate the ΔZF10 RNA-binding mutant in this study, as our focus was on the role of CTCF-RNA interactions. The intact CTCF-cohesin interactions in the ΔZF1 mutant observed by ChIP-MS is consistent with results from CTCF-IP and Rad21-immunoblots published previously^41^.

Although a gross loss in CTCF-cohesin interactions was undetectable, we considered that there might be changes in CTCF-cohesin colocalization at CTCF anchors in the ΔZF1 mutant. Thus, we performed ChIP-seq of Rad21 and Smc3, two of the cohesin core subunits. There were minimal to no changes in Rad21 or Smc3 ChIP-seq peaks in ΔZF1 (**Fig. S6C, S6D, S6E, S6F**). We also found no differences in Rad21 or Smc3 ChIP-seq enrichment between WT and ΔZF1 at both ΔZF1-lost and ΔZF1-retained anchors (**Fig. S6G, S6H, Fig. S6I, S6J**). **Fig. 3F** shows a representative *Podxl* locus wherein an NPC-specific chromatin loop is lost in NPC-ΔZF1, while CTCF, Rad21, and Smc3 DNA-binding remain unchanged at the anchors. Based on these results, we concluded that the loss of CTCF loop anchors in the ΔZF1 mutant was not due to a loss of CTCF-DNA binding nor of CTCF-cohesin interactions and instead, was most likely due to the loss of direct CTCF-RNA interactions.

### Dysregulated genes in the ΔZF1 mutant are enriched at disrupted loops

To assess the effect of the ΔZF1 mutant on gene expression in ESCs and NPCs, we performed bulk RNA-seq followed by DESeq2 analysis^62^. We compared gene expression from ESC-ΔZF1 and NPC-ΔZF1 to their WT counterparts. In ESC-ΔZF1, we identified 432 dysregulated genes, with 235 downregulated and 197 upregulated (**Fig. S7A, Table S3**). In NPC-ΔZF1, we found 2,060 dysregulated genes, with 1,226 downregulated and 834 upregulated (**Fig. 4A, Table S4**). Given that the change in gene expression was greater in the case of NPCs, we asked whether CTCF-ZF1 is crucial for an NPC-specific transcriptional program. We performed a GO term enrichment analysis using PANTHER^63^, and found that the dysregulated genes in NPC-ΔZF1 had an overrepresentation of terms related to NPC function, such as dendrite morphogenesis and development, regulation of neurogenesis, and nervous system development (**Fig. 4B, Table S5**).

**Figure 4.**
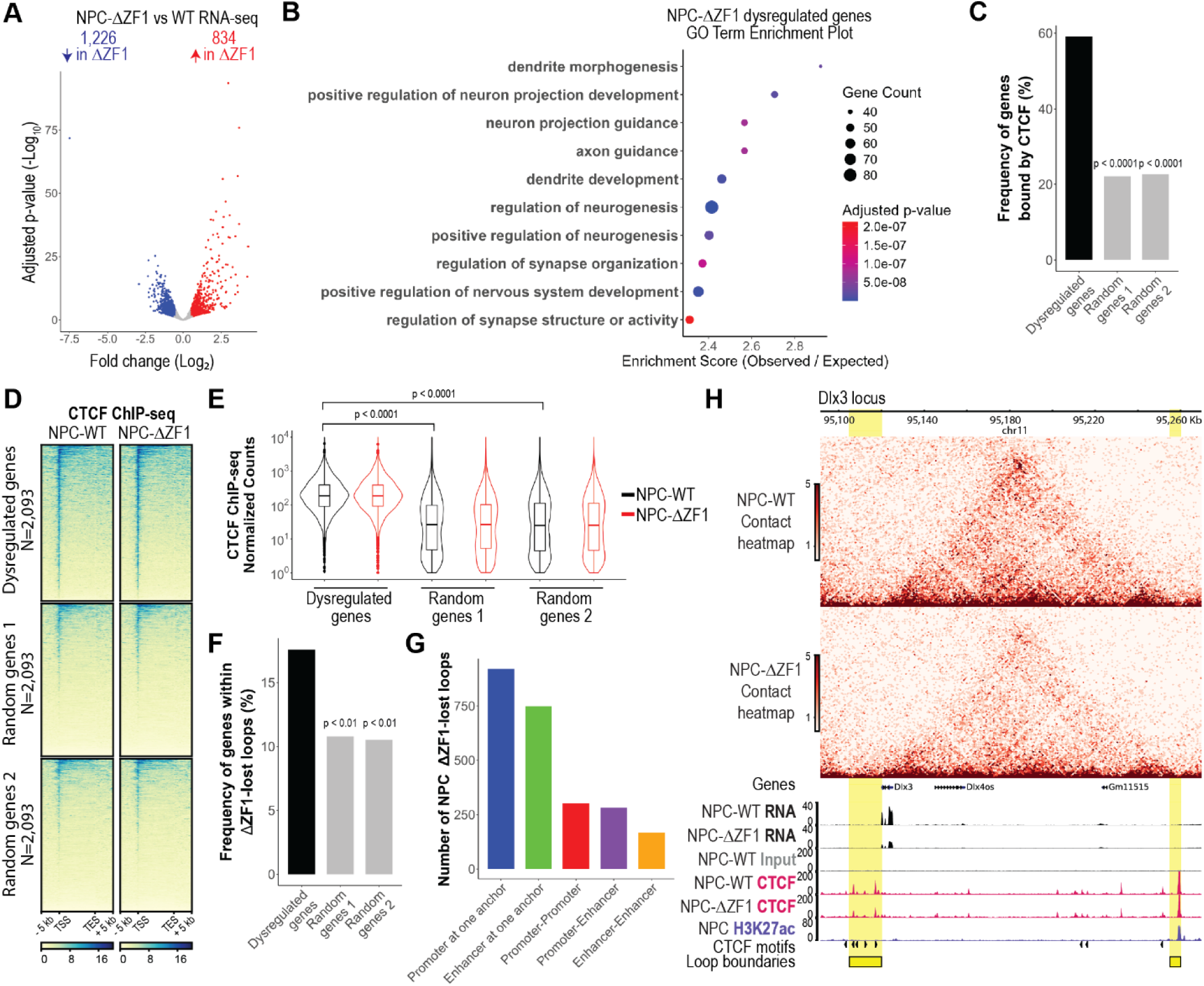
Dysregulated genes in NPC-ΔZF1 mutant are enriched at disrupted loops. (A) Deseq2 volcano plots showing gene expression changes in NPC-ΔZF1 compared to NPC-WT (see all in **Table S4**). Adjusted p-value cutoff: ≤ 0.05, log_2_ fold-change cutoff: ≥ 0.5, ≤-0.5, N=2. (B) Panther GO term analysis of NPC-ΔZF1 dysregulated genes (see all in **Table S5**). Shown are the top 10 terms related to neuronal function. (C) Bar plots showing the percentage of genes from each gene set (dysregulated, random genes 1, random genes 2) that overlap with CTCF ChIP-seq peaks in NPCs. p-values comparing each of the random gene sets to dysregulated genes were determined using Fisher’s Exact test. (D) CTCF ChIP-seq heatmaps at the TSS to TES of dysregulated genes and two sets of randomly generated genes in NPCs. (E) Quantification of CTCF ChIP-seq reads from TSS to TES of genes shown in (D). p-values were determined using t-test. (F) Bar plots showing the percentage of genes from each gene set (dysregulated, random genes 1, random genes 2) that are co-localized within ΔZF1-lost loops in NPCs. p-values comparing each of the random gene sets to dysregulated genes were determined using Fisher’s Exact test. (F) Bar plots showing the number of NPC-ΔZF1-lost anchors that overlapped with promoters and/or enhancers. (G) An example locus with decreased enhancer-promoter loop contact in ΔZF1 mutant. Micro-C contact heatmap (above) compares WT and ΔZF1 contact frequency at the loop anchors (highlighted in yellow). *Dlx3* gene expression is shown in the RNA-seq tracks below. CTCF ChIP-seq tracks show CTCF-binding at the loop anchors, wherein the upstream anchor is at the *Dlx3* promoter and the downstream anchor is at a putative enhancer marked by H3K27ac. H3K27ac ChIP-seq is reanalyzed data from Tiwari et al., 2018 (see **Table S6**). The presence of CTCF motifs and their orientation are annotated.

To assess whether gene expression changes were directly regulated by CTCF, we examined whether there was an enrichment of CTCF binding at dysregulated genes compared to randomly generated lists of genes. We found that 63% of ESC-ΔZF1 dysregulated genes (**Fig. S7B**) and 59% of NPC-ΔZF1 dysregulated genes (**Fig. 4C**) were bound by CTCF, compared to 22-24% of random genes (**Fig. 4C, S7B**). Indeed, CTCF ChIP-seq heatmaps revealed that dysregulated genes had a significant enrichment in CTCF binding, particularly at TSSs (**Fig. 4D, Fig. S7C**). We quantified the CTCF ChIP-seq read counts across the genes [from TSS to transcription end site (TES)] and found that CTCF binding was significantly more enriched at ΔZF1-dysregulated genes compared to random genes, with no changes in CTCF binding in the ΔZF1 mutant (**Fig. 4E, Fig. S7D**).

We further investigated whether dysregulated genes were co-localized within ΔZF1-lost anchors. Our analysis revealed that 18% (80 out of 436) of dysregulated genes in ESC-ΔZF1 (**Fig. S7E**) and 18% (368 out of 2,093) of dysregulated genes in NPC-ΔZF1 were co-localized at disrupted CTCF anchors, compared to 10-11% of random genes (**Fig. 4F**). Statistical analysis indicates that dysregulated genes were significantly more likely to co-localize within ΔZF1-lost loops than would be expected by random chance (**Fig. 4F, S7E**).

Next, we annotated the loop anchors based on their overlap with promoters or putative enhancers. Putative enhancers were identified based on ChIP-seq peaks of histone H3 acetylated at lysine 27 (H3K27ac) in ESCs and NPCs from previous studies (**Table S6**)^64,65^. We found several hundred ΔZF1-lost anchors overlapped with either a promoter (P) or enhancer (E) at one anchor, or showed P-P, P-E, and E-E overlaps at both anchors (**Fig. 4G, Fig. S7F**). We show an example of a ΔZF1-lost anchor wherein the upstream anchor overlaps with the promoter of a ΔZF1-downregulated gene, *Dlx3* (**Fig. 4H**). In the ΔZF1 mutant, the CTCF-bound *Dlx3* promoter exhibited decreased looping contact with a downstream putative enhancer marked by H3K27ac, likely leading to decreased *Dlx3* expression (**Fig. 4H**).

Although some genes may be dysregulated indirectly due to a transcriptional cascade rather than direct chromatin loop changes, the higher enrichment of CTCF binding at dysregulated genes and the increased likelihood of dysregulated gene co-localization at ΔZF1-lost anchors suggest that disruptions in chromatin architecture directly contributed to transcriptional changes in the ΔZF1 mutant. These transcriptional changes can be partially explained by the disruption of CTCF-bound promoter-enhancer contacts.

### Identification of NPC-specific CTCF-ZF1 RNA interactions

In light of our findings thus far, we hypothesized that RNAs upregulated in NPCs interact with CTCF to facilitate the formation of NPC-specific CTCF anchors *in cis*, *i.e.* at or near the site of transcription. To identify potential loop-inducing CTCF-RNA interactions, we performed CLIP-seq (iCLIP2)^66^. We used anti-Flag immunoprecipitation to purify Flag-Halo-tagged rescue CTCF (WT or ΔZF1) and verified CTCF-RNA interactions with RNA-^32^P labeling and CTCF immunoblotting (**Fig. S8A**). We then sequenced the CTCF-RNA interactome in ESCs and NPCs. We used PureCLIP to identify CTCF-RNA single-nucleotide crosslinks^67^ and did post-processing of PureCLIP-called crosslinks^68^ to call reproducible CLIP peaks. In ESCs, we identified 69,685 peaks in CTCF-WT, whereas CTCF-ΔZF1 had 33,354 peaks (**Fig. 5A**). In NPCs, there were 50,758 peaks in CTCF-WT, while CTCF-ΔZF1 had 30,799 peaks (**Fig. 5B**). CLIP-seq heatmaps of called peaks showed a higher enrichment of CTCF-RNA crosslinks in WT compared to ΔZF1 mutant in both ESCs (**Fig. 5C, 5D**) and NPCs (**Fig. 5E, 5F**). WT CLIP peaks were annotated to 6,438 genes in ESCs and 6,350 genes in NPCs. Accordingly, CLIP-seq enrichment across the TSSs to TESs of annotated genes were decreased in ΔZF1 (**Fig. S8B, S8C**). CLIP peaks were mapped to multiple RNA features, including 5’UTRs, 3’UTRs, exons, and introns (**Fig. S8D**). Annotated genes ranged from coding, non-coding, antisense, to pseudogenes, with the majority being coding RNAs (**Fig. S8E**). *De novo* RNA motif analysis using MEME^69^ revealed significant RNA-binding motifs, though they were found only in ∼10-20% of CLIP peaks (**Fig. S8F**).

**Figure 5.**
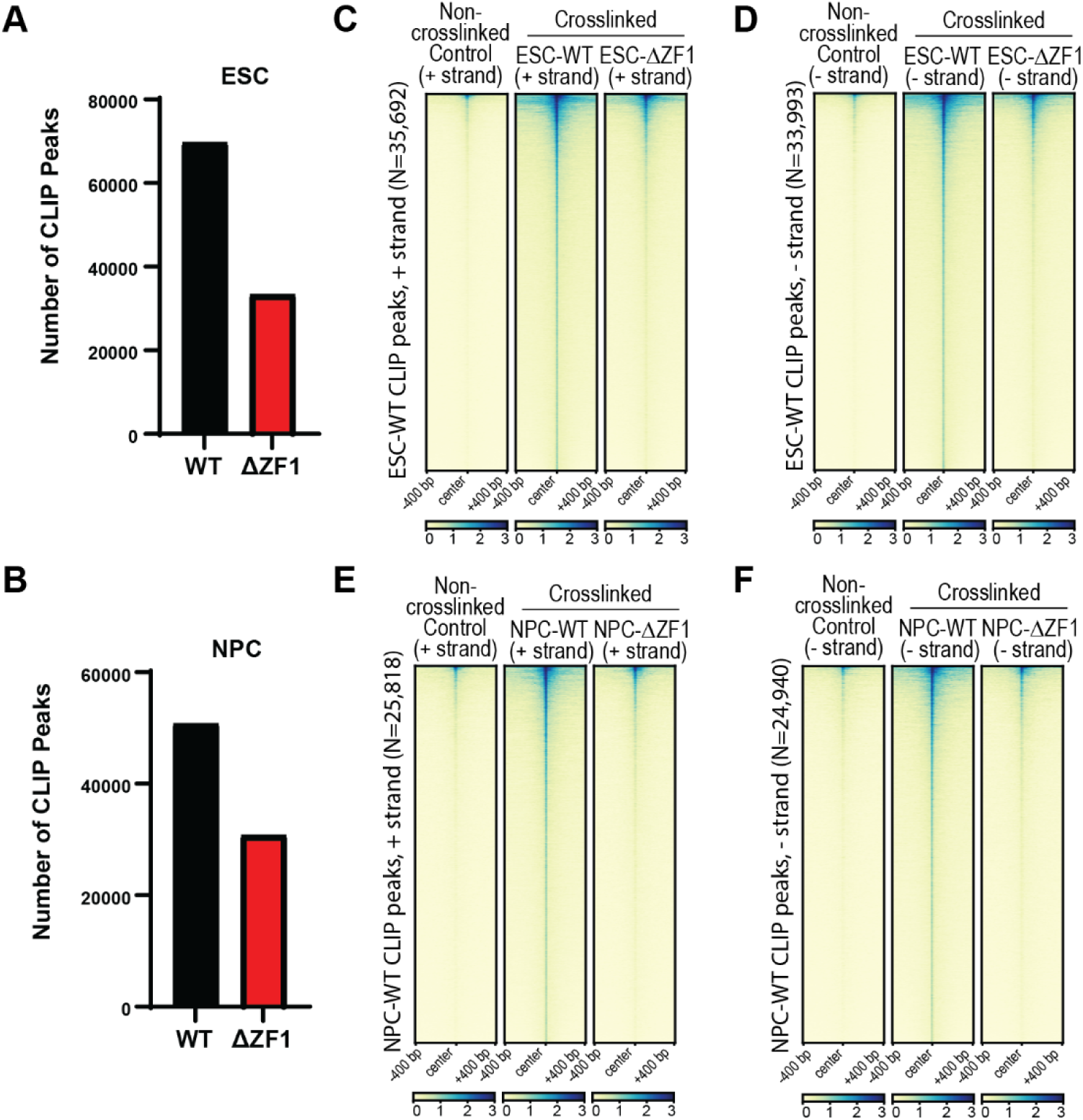
Decreased CTCF-RNA crosslinking in the ΔZF1 mutant. (A, B) ESCs (A) and NPCs (B) were UV-crosslinked and Flag-halo tagged WT or ΔZF1 rescue CTCF was immunoprecipitated using anti-Flag beads. RNA was purified and cDNA library was sequenced. Reproducible CLIP peaks were called (see Methods) in WT and ΔZF1 mutant. Bar graphs of the number of reproducible peaks are shown (N=3-4). (C, D) Heatmaps of ESC-CTCF CLIP peak enrichment in the non-crosslinking negative control, and crosslinked WT and ΔZF1. CLIP peaks are shown for the (C) plus (+) and (D) minus (-) strands. (E, F) Heatmaps of NPC-CTCF CLIP peak enrichment in the non-crosslinking negative control, and crosslinked WT and ΔZF1. CLIP peaks are shown for the (E) plus (+) and (F) minus (-) strands.

To identify NPC-specific CTCF-RNA interactions, we used DESeq2 to call genes with significantly higher CLIP-seq reads in NPC WT compared to ESC WT. This analysis revealed 554 NPC-specific CLIP-seq genes (**Fig. 6A, Table S7**). Not surprisingly, RNA-seq data showed that the majority of these genes were upregulated in NPCs compared to ESCs (**Fig. 6B**). Additionally, most of the NPC-specific genes did not exhibit significantly decreased expression in the NPC-ΔZF1 mutant compared to NPC-WT (**Fig. 6B**), suggesting that the overall reduction in CTCF-RNA interactions in the NPC-ΔZF1 mutant was not due to decreased RNA expression. To narrow down the RNA candidates for functional validation, we overlapped the list of 554 NPC-specific genes with that of genes that were upregulated in NPCs (relative to ESCs) and the list of genes co-localized at NPC-specific CTCF anchors (**Fig. 6C**). Among these RNAs, we selected *Podxl* and *Grb10* for further study, as their loci are associated with NPC-specific, ΔZF1-lost loops (**Fig. 2E** and **Fig. 2F**, respectively), they are expressed in the mouse brain and have known functions in neurons^70–72^. Closer inspection of the RNA-seq and CLIP-seq tracks of these genes showed that they are upregulated in NPCs and have decreased interaction with CTCF in NPC-ΔZF1 despite their expression levels being unaffected (**Fig. 6D, 6E**).

**Figure 6.**
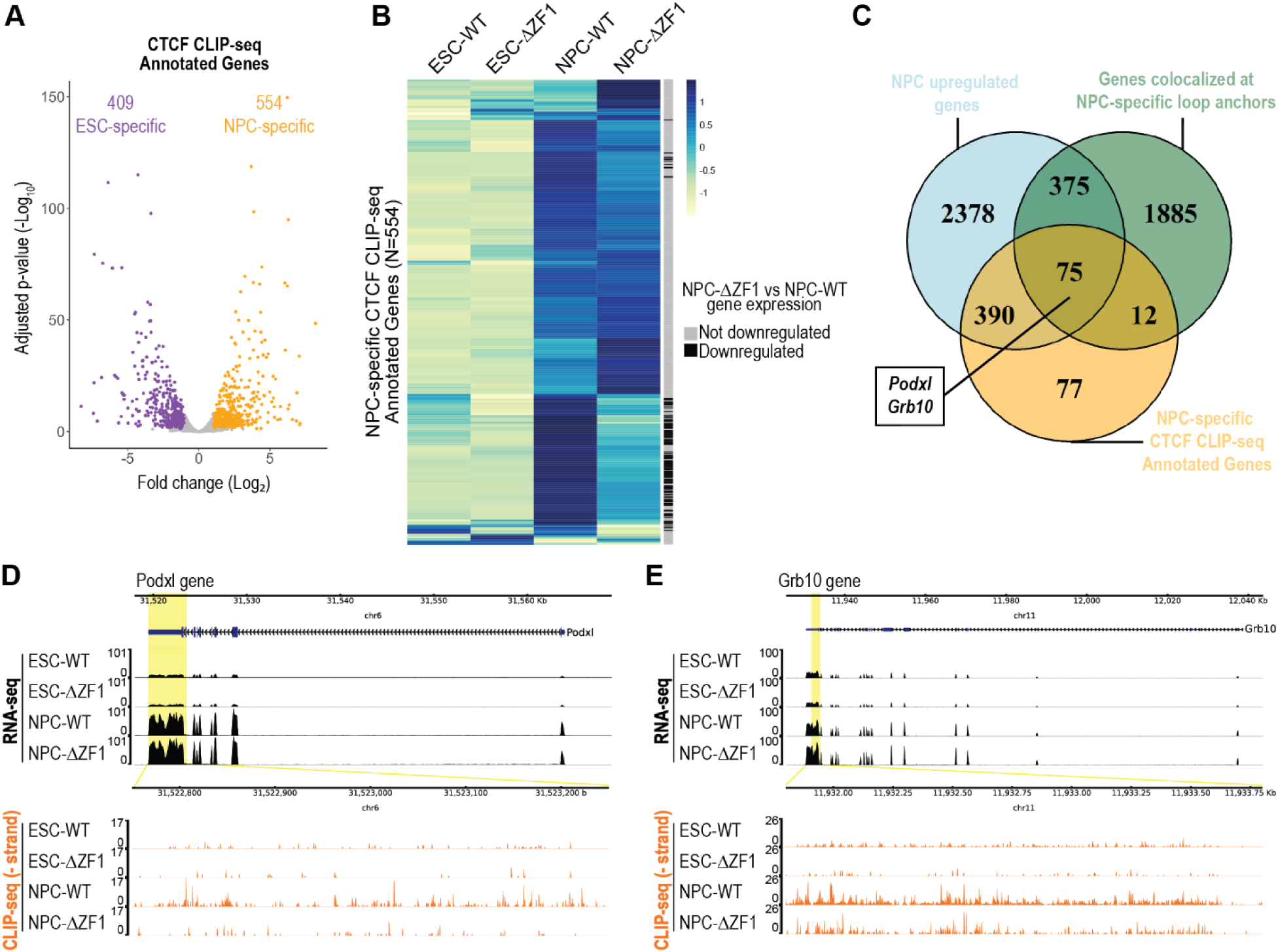
Identification of NPC-specific CTCF-RNA interactions. (A) Deseq2 volcano plot showing differential CTCF-RNA interactions comparing NPCs vs ESCs (see all in **Table S7**). Each dot is a gene. Total CLIP-seq read counts from the TSS to TES of genes were compared. Adjusted p-value cutoff: ≤ 0.05, log_2_ fold-change cutoff: ≥ 1, ≤-1, N=3-4. (B) RNA-seq heatmaps of NPC-specific, CLIP-seq annotated genes, comparing ESC-WT, ESC-ΔZF1, NPC-WT, and NPC-ΔZF1. Each row is a gene and is annotated based on whether or not gene expression in NPC-ΔZF1 is significantly downregulated compared to WT. (C) Venn diagram of i) Genes upregulated in NPCs relative to ESCs, ii) genes colocalized at NPC-specific loop anchors, and iii) genes annotated to be NPC-specific, CTCF-interactors by CLIP-seq. Genes at the intersection of the three gene lists were considered for functional validation with *Podxl* and *Grb10* being selected. (D, E) RNA-seq tracks (above) and CLIP-seq tracks (below) at genes of interest: (D) *Podxl* and (E) *Grb10*. CLIP-seq tracks are zoomed in for better visualization of nucleotide-level crosslinks.

### Truncation of NPC-specific RNAs interacting with CTCF disrupts NPC-specific loops *in cis*

Transcriptional upregulation of genes after differentiation into NPCs may facilitate chromatin looping through their RNA products interacting with CTCF at nearby CTCF binding sites. We hypothesized that truncation of the candidate RNAs, *Podxl* and *Grb10*, would decrease CTCF anchor enrichment *in cis*, specifically at the CTCF anchors where one anchor overlaps with the transcribed gene. To test this hypothesis, we disrupted the transcription of *Podxl* and *Grb10* RNAs independently by inserting T2A-eGFP followed by an SV40-polyA (pA) transcription termination signal downstream of the first exon of the coding sequence of each gene (**Fig. 7A**). Using CRISPR-Cas9 gene editing, we generated homozygous Podxl-T2A-eGFP-pA (Podxl pA) and Grb10-T2A-eGFP-pA (Grb10 pA) RNA truncation mutants (**Fig. 7A**). The truncation of *Podxl* and *Grb10* was verified by RT-qPCR (**Fig. S9A, S9B**) and RNA-seq counts (**Fig. 7B, 7C**). The mutant lines were eGFP fluorescent, resulting from in-frame knock-in of the repair template DNA (**Fig. S9C**). Both pA mutants were able to differentiate into NPCs expressing the Sox1 marker (**Fig. S9D**). Importantly, the mutants did not show changes in CTCF or Rad21 protein levels (**Fig. S9E**).

**Figure 7.**
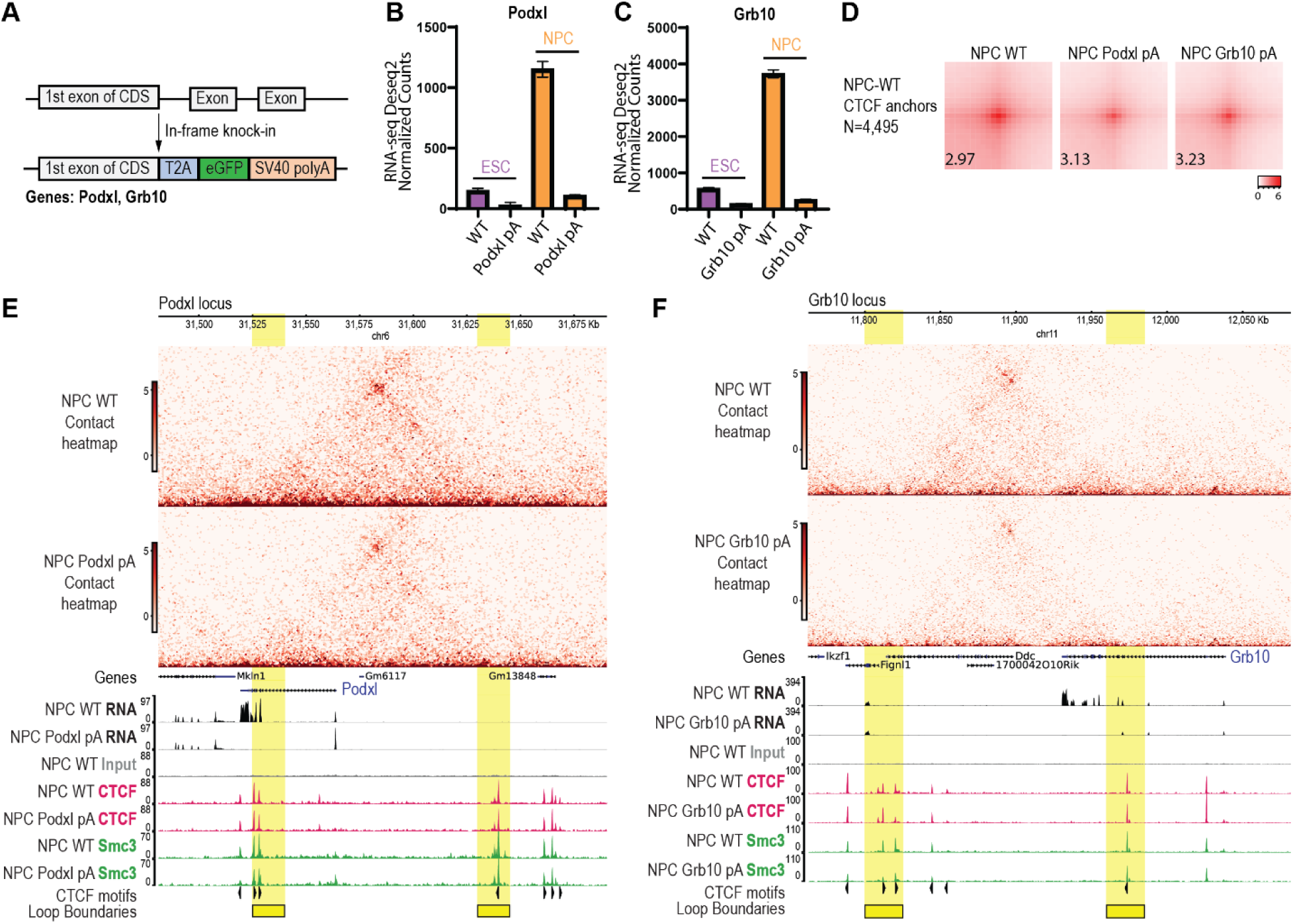
Truncation of NPC-specific RNAs interacting with CTCF leads to decreased chromatin loops *in cis*. (A) Schematic of CRISPR-Cas9-mediated truncation of target RNAs, *Podxl* and *Grb10*. We generated a homozygous in-frame knock-in of T2A-eGFP-SV40 polyA (pA) downstream of the 1^st^ exon of the coding sequence (CDS) of each gene. (B, C) RNA-seq Deseq2 normalized read counts of (B) *Podxl* in WT and Podxl pA mutant and (C) *Grb10* in WT and Grb10 pA mutant. Data are represented as mean ± SEM, N=3. (D) APA plots of NPC WT CTCF anchors, comparing WT, NPC Podxl pA, and NPC Grb10 pA. Numbers indicate APA score. (E, F) Micro-C contact heatmaps (above) and RNA-seq tracks and ChIP-seq tracks (CTCF and Smc3) (below) at (E) the *Podxl* locus or (F) the *Grb10* locus. NPC WT is compared to either the Podxl pA mutant or the Grb10 pA mutant, respectively. Loop boundaries are highlighted in yellow. The presence of CTCF motifs and their orientation are annotated.

We performed Micro-C on the parental NPC WT cells, as well as NPC Podxl pA and Grb10 pA mutants, with higher read depth than previously used for the CTCF rescue lines (**Table S8**) to resolve loops more clearly at the *Podxl* and *Grb10* loci. Unlike the NPC-ΔZF1 mutant, we found that previously identified NPC-WT rescue CTCF anchors were overall not affected in either Podxl pA or Grb10 pA NPCs (**Fig. 7D**). However, the CTCF anchors colocalized at the *Podxl* and *Grb10* genes were now decreased in the Podxl pA and Grb10 pA mutants, respectively (**Fig. 7E, 7F**), similar to the case of NPC-ΔZF1. Yet, there were no changes in CTCF and Smc3 DNA-binding at these loci (**Fig. 7E, 7F**). The more widespread loss of loops in the NPC-ΔZF1 compared to single RNA truncations suggests that multiple RNAs interact with CTCF-ZF1 to maintain loops genome-wide.

In conclusion, the RNA-binding region of CTCF, specifically the ZF1 region, is critical for maintaining NPC-specific chromatin loops at the *Podxl* and *Grb10* locus, among others. CTCF-ZF1 interacts with NPC-upregulated RNAs, such as *Podxl* and *Grb10*. Disruption of CTCF-RNA interactions through either the deletion of ZF1 or the truncation of *Podxl* or *Grb10* RNA led to decreased chromatin loop enrichment *in cis*.

## DISCUSSION

CTCF is a key regulator of chromatin loops^17^, but the mechanisms underlying how CTCF facilitates chromatin loops with cell-type-specificity have remained unclear. Our data suggest that cell-type-specific RNAs orchestrate chromatin loops *in cis* through interactions with CTCF, adding a new dimension to our understanding of chromatin architecture during differentiation. The implications of our findings extend beyond the differentiation of ESCs into NPCs, as the principles uncovered in this study are likely relevant in other differentiation pathways. For example, CTCF is crucial in forming chromatin loops that drive myogenic transdifferentiation^73^, cardiomyocyte development^74^, and pancreatic cell differentiation^75^. Furthermore, CTCF-RNA-mediated loops may provide a positive feedback mechanism for transcription. In this scenario, cell-type-specific transcription factors initiate gene upregulation, leading to CTCF-RNA-mediated promoter-enhancer looping, which in turn reinforces gene upregulation.

The precise mechanisms by which CTCF-ZF1 RNA interactions facilitate chromatin loop formation during differentiation remain to be elucidated. Although we found some reduction in the levels of CTCF bound to its cognate sites in the ΔZF1 mutant, this reduction alone is insufficient to explain the far more pronounced loss of chromatin loops observed. Thus, the loss of CTCF anchors in ΔZF1 is not due to the loss of CTCF chromatin recruitment. In accordance, the truncation of CTCF-interacting RNAs, *Podxl* and *Grb10*, did not lead to decreased CTCF chromatin binding, despite an observable decrease in chromatin looping. Thus, the mechanism fostering CTCF loop anchor formation by ZF1-mediated RNA interaction appears distinctive from CTCF-mediated loop formation involving other, previously described non-coding RNAs (ncRNA) and enhancer RNAs (eRNAs) given their primary role in either recruiting or releasing CTCF from chromatin^44–50^. For example, long non-coding RNA (lncRNA), *Jpx*, binds to and extricates CTCF from low-affinity binding sites^46,48^. Consequently, the loss of *Jpx* in ESCs leads to increased CTCF DNA-binding and chromatin looping^48^. Conversely, ncRNA *MYCNOS* was shown to be important in recruiting CTCF to the MYCN promoter in neuroblastoma cell lines^47^, while eRNAs recruit CTCF to the boundary of the INK4a/ARF TAD in HeLa cells^49^. Similarly, ncRNAs *Tsix* and *Xite* recruit CTCF to the X inactivation center^44^. Whether the regulation of CTCF chromatin binding by these RNAs relies on known CTCF RNA-binding regions (ZF1, ZF10, and/or RBRi) has not been investigated.

Although the truncation of *Podxl* and *Grb10* RNAs would also lead to a loss of their protein products, Podxl and Grb10 are mainly plasma membrane^72^ and cytoplasmic^76^ proteins, respectively, and do not directly interact with CTCF, based on CTCF ChIP-MS results (not shown). Hence, it is highly likely that the RNA molecules themselves, rather than their protein counterparts, are crucial for the direct maintenance of chromatin loops. Indeed, our study uncovered coding RNAs as being the predominant RNA biotype in our CTCF CLIP-seq. This makes the regulatory landscape of the genome more complex than previously envisioned, with coding RNAs playing multifaceted roles that extend beyond their traditional functions. Future studies could explore how widespread this phenomenon is and whether other coding RNAs have similar structural roles in different cellular contexts.

Interestingly, our study also suggests that the loss of CTCF loop anchors in the ΔZF1 mutant is not due to a loss of cohesin interaction nor of CTCF-cohesin colocalization on chromatin. This was a surprising finding given the complete loss of chromatin loops upon cohesin degradation^21^. It is possible that loop extrusion by cohesin initially brings CTCF anchors together and that CTCF-RNA interactions maintain the loop even after cohesin is unloaded, thus increasing loop lifetime. Therefore, CTCF-RNA interactions can independently contribute to chromatin loop maintenance, in addition to the well-established role of cohesin in loop extrusion. One possibility is that CTCF forms dimers or higher-order oligomers mediated by RNA interactions, stabilizing the loop structure, as initially suggested in previous studies^40,41,43^. Alternatively, RNA molecules may act as scaffolds or bridges that bring distant CTCF-bound chromatin regions into proximity, thereby promoting loop formation.

An intriguing and important aspect of our findings is that the CTCF-bound loops that were lost upon RNA truncation are in close proximity to the site of transcription. This putative *cis*-acting mechanism could be crucial in maintaining the integrity of chromatin loops in a localized or specific manner. The dual requirement of transcription and CTCF proximity could create an efficient system for dynamically regulating chromatin loops in response to cellular signals and differentiation cues. Future studies to explore how RNA proximity influences CTCF-mediated loop formation could entail artificial tethering of RNAs to distant sites or altering the site of transcription to assess its impact on loop stability.

The RNA motifs or secondary structures that mediate CTCF-RNA interactions are of significant interest. Previous CTCF-RNA EMSAs demonstrated that CTCF exhibits higher affinity for certain RNAs over others^44^, suggesting specificity in these interactions. Our study supports this notion, as not all NPC-upregulated RNAs interacted with CTCF, indicating selective binding. In the absence of a prevalent RNA recognition motif, specificity is likely driven by RNA secondary or tertiary structures recognized by the ZF1 region. Elucidating the structural mechanisms by which CTCF maintains chromatin loops in conjunction with RNA remains a challenging, yet promising and tenable research direction.

In addition to interacting with CTCF, RNA molecules are known to modulate the function of other chromatin-associated proteins, such as Polycomb Repressive Complex 2 (PRC2). A key regulator of gene silencing and chromatin compaction, PRC2, catalyzes histone H3 trimethylation at lysine 27 (H3K27me3). Its activity is influenced by RNA. For instance, nascent and G-quadruplex RNAs interact with PRC2 and inhibit its methyltransferase activity^77–80^. RNA has also been shown to promote PRC2 chromatin occupancy^81^. In contrast, other studies suggest that RNAs facilitate H3K27 trimethylation^82^ and compete with chromatin for PRC2 binding^83^. Thus, the function of PRC2-RNA interactions has been a subject of significant debate and controversy. Despite these differing viewpoints, PRC2-RNA interactions underscore that there are potential, multifaceted roles of RNA in chromatin regulation, extending beyond their involvement with CTCF. The ability of RNA to interact with multiple chromatin-associated proteins suggests a complex network of RNA-mediated regulatory mechanisms that coordinate chromatin architecture and gene expression.

Lastly, future explorations of the mechanisms by which CTCF-RNA interactions contribute to chromatin architecture may be impactful in the context of neuronal pathologies. CTCF is crucial for NPC proliferation, differentiation, and survival,^84^ and CTCF mutations are linked to neurological disorders such as intellectual disability^85^, autism spectrum disorders^86^, and neurodegenerative diseases^87^. Our findings highlight the importance of CTCF-RNA interactions in maintaining the chromatin structure of NPCs. By facilitating the formation of NPC-specific chromatin loops, CTCF-RNA interactions ensure the proper regulation of gene expression necessary for NPC function and differentiation. Understanding these interactions in greater detail could provide new insights into the molecular basis of neuronal function, in both health and disease.

### Limitations of the Study

Our study provides significant insight into the role of CTCF-RNA interactions in chromatin loop maintenance in the context of cellular differentiation. However, some limitations should be acknowledged. First, our findings are based on in vitro models of ESC to NPC differentiation, which may not fully recapitulate the complexity of in vivo differentiation processes. We also did not disentangle the functions of the RNA molecules from their protein products. While truncation of *Podxl* and *Grb10* RNAs led to decreased chromatin loop enrichment, it remains unclear whether the observed effects are solely due to the loss of RNA interactions with CTCF or if the absence of the corresponding proteins also played an indirect role. Importantly, while we have demonstrated that CTCF-RNA interactions are required for maintaining chromatin loops, we have not demonstrated that these interactions are sufficient to establish or maintain these loops. Finally, while our study focused on two RNA candidates, future research may systematically identify and validate additional RNAs involved in CTCF-mediated chromatin architecture across different cell types or developmental stages.

## RESOURCE AVAILABILITY

### Lead contact

Further information and requests for resources and reagents should be directed to and will be fulfilled by the lead contact, Danny Reinberg.

### Materials availability

The plasmids generated in this study are deposited to Addgene. All unique/stable materials generated in this study are available from the corresponding author upon reasonable request with a completed Materials Transfer Agreement.

### Data and code availability

- Sequencing data has been deposited at Gene Expression Omnibus (GEO), accession numbers GSE287375, GSE287376, GSE287379, GSE287380. Publicly available datasets used are listed in Table S6. Raw data of images used in Figures 1B, S1C, S8A, S9E were deposited in Mendeley (doi: 10.17632/2mcy77yh5y.1).
- This paper does not report original code.
- Any additional information required to reanalyze the data reported in this paper is available from the lead contact upon request.

## ACKNOWLEDGMENTS

We thank L. Vales for comments and advice on the manuscript; J. Skok, A. Tsirigos, and E. Nudler for feedback as the work was in progress; past and present members of the Reinberg and Mazzoni laboratories for discussions; D. Hernandez and D. Beck for cryostorage space. We also thank the NYU Grossman School of Medicine’s Genome Technology Center, particularly P. Zappile and E. Grasso for sequencing services; Rutgers University Mass Spectrometry Facility, particularly H. Zheng for conducting LC-MS/MS procedure; and staff at NYU Grossman School of Medicine’s Cytometry and Cell Sorting Core Facility. This study utilized computing resources at the High-Performance Computing Facility of the Center for Health Informatics and Bioinformatics at the NYU Grossman School of Medicine. Funding: This work was supported by the National Institutes of Health (NIH) grant R01NS100897, National Cancer Institute (NCI) grant 5R01CA199652-21, and the Howard Hughes Medical Institute (D.R.), and NIH grant R01NS100897 (E.O.M.). The NYU Grossman School of Medicine’s Genome Technology Center and the NYU Grossman School of Medicine’s Cytometry and Cell Sorting Core are supported partially by the NIH/NCI Support Grant P30CA016087 at the Laura and Isaac Perlmutter Cancer Center.

## AUTHOR CONTRIBUTIONS

K.L., P.Y.H., and D.R. conceived the project and designed the experiments. K.L. and D.R. wrote the paper. K.L., P.Y.H., and X.Q. generated the cell lines. K.L. performed the experiments. K.L. and S.H. performed the bioinformatic analyses. E.O.M. advised on the progression of this study.

## SUPPLEMENTARY TABLES

**Table S1-S10.**

**Table S1.** ChIP-MS analysis of ESC-ZF1 mutant vs ESC-WT, related to Figure S6A (Separate Excel file)

**Table S2.** ChIP-MS analysis of ESC-ZF10 mutant vs ESC-WT, related to Figure S6B (Separate Excel file)

**Table S3.** RNA-seq analysis of ESC-ZF1 mutant vs ESC-WT, related to Figure S7A (Separate Excel file)

**Table S4.** RNA-seq analysis of NPC-ZF1 mutant vs NPC-WT, related to Figure 4A (Separate Excel file)

**Table S5.** Panther GO Term Enrichment of dysregulated genes in NPC-ZF1 mutant, related to Figure 4B

(Separate Excel file)

**Table S6.** Public datasets used in this study, related to Figures 4G, 4H, S7F.

| Cell line | Assay type | Target | Database | Experiment ID |
| --- | --- | --- | --- | --- |
| *Mus musculus* strain 129/Ola ES-E14 | ChIP-seq | H3K27ac | ENCODE | doi:10.17989/ENCSR000CGQ |
| *Mus musculus* strain C57/BL-6 ES-derived neural progenitor cells | ChIP-seq | H3K27ac | NCBI | GSM2535239 |
| *Mus musculus* strain C57/BL-6 ES-derived neural progenitor cells | ChIP-seq | H3K27ac | NCBI | GSM2535240 |

**Table S7.** CLIP-seq annotated genes in NPCs vs ESCs, related to Figure 6A (Separate Excel file)

**Table S8.** Micro-C read depth, related to Figures 2, 7, S3, S4, S5, and Methods (Separate Excel file)

**Table S9.** Oligonucleotides used in this study, related to Figure 1A, S1B, S2A, S9A, S9B, and Methods

(Separate Excel file)

**Table S10.** Antibodies used in this study, related to Figures 1B, 1C, 3, S1C, S5, S6, S8A, S9D, S9E, and Methods.

| Primary antibodies | Company | Catalogue # | Application | Dilutions |
| --- | --- | --- | --- | --- |
| Rabbit Anti-CTCF | Abcam | ab70303 | ChIP, WB | ChIP: 2 ug/300 ug chromatin, WB: 1:5,000 |
| Mouse Anti-Flag | Sigma | F1804 | WB | 1:500 |
| Rabbit Anti-Lamin B1 | Abcam | ab16048 | WB | 1:20,000 |
| Goat Anti-Sox1 | R&D Systems | AF3369 | Flow cytometry | 1:200 from 500 ug/mL reconstituted stock |
| Rabbit Anti-Rad21 | Abcam | ab217678 | ChIP, WB | ChIP: 4 ug/300 ug chromatin, WB: 1:5,000 |
| Rabbit Anti-Smc3 | Abcam | ab9263 | ChIP | 2 ug/300 ug chromatin |
| Mouse Anti-Oct3/4 | Santa Cruz | sc-5279 | WB | 1:10,000 |
| **Secondary antibodies** | **Company** | **Catalogue #** | **Application** | **Dilutions** |
| Donkey Anti-Goat IgG (H+L) Highly Cross-Adsorbed Secondary Antibody, Alexa Fluor™ Plus 647 | Invitrogen | A32849 | Flow cytometry | 1:1,000 |
| Goat Anti-Rabbit IgG (H+L)-HRP Conjugate | Bio-Rad | 1706515 | WB | 1:10,000 |
| Goat Anti-Mouse IgG (H+L)-HRP Conjugate | Bio-Rad | 1706516 | WB | 1:5,000 |

## STAR★METHODS

## KEY RESOURCES TABLE

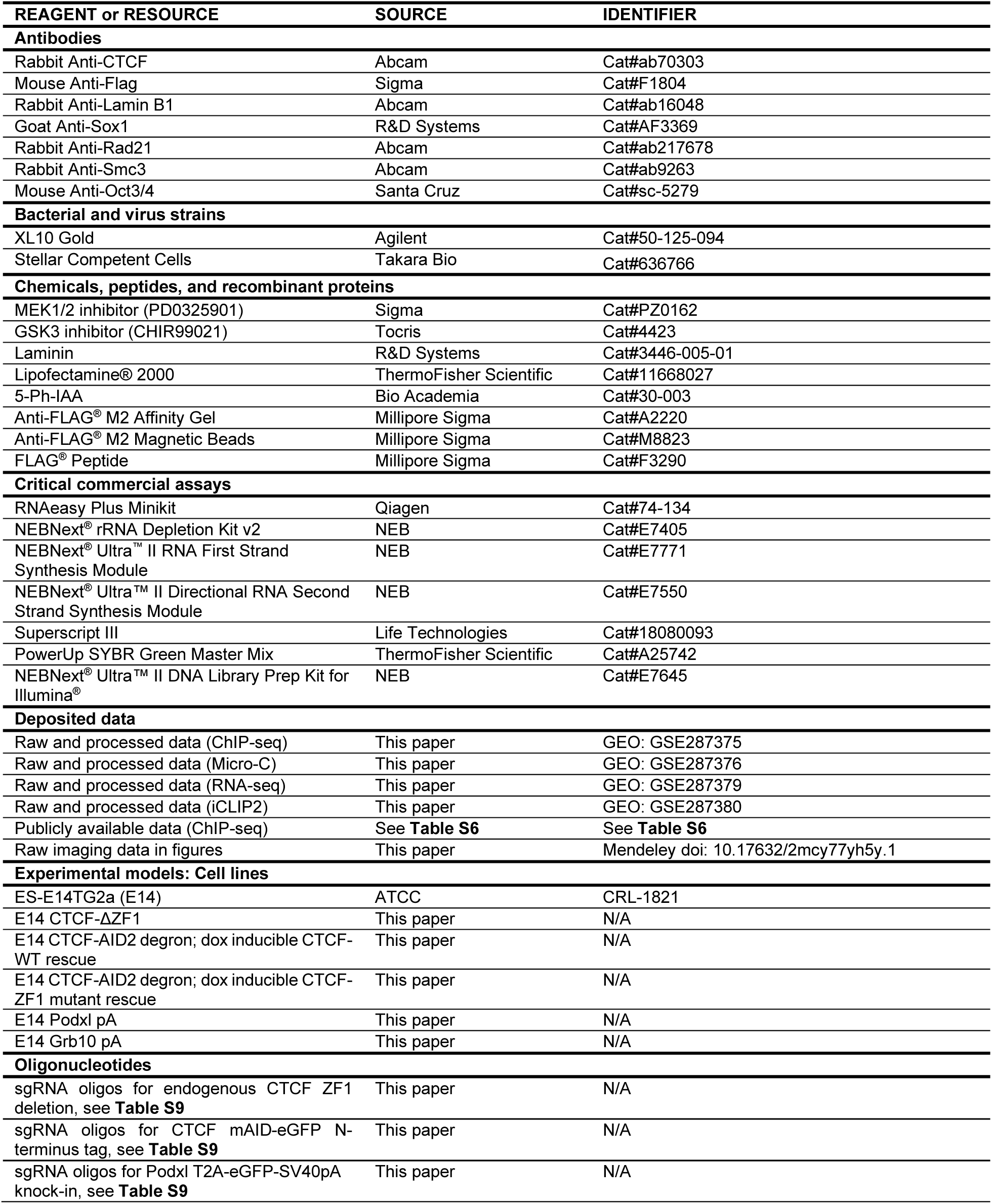
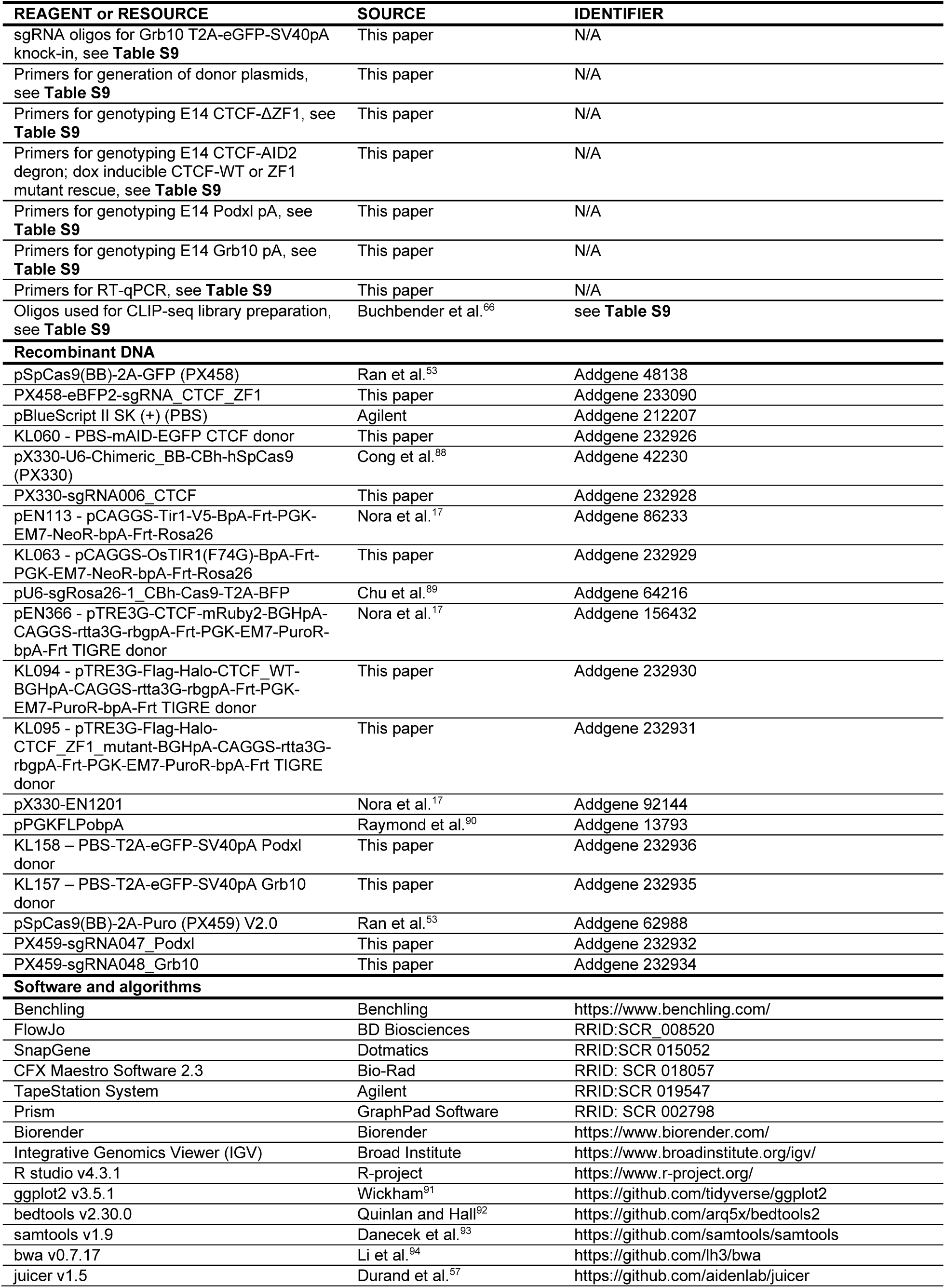
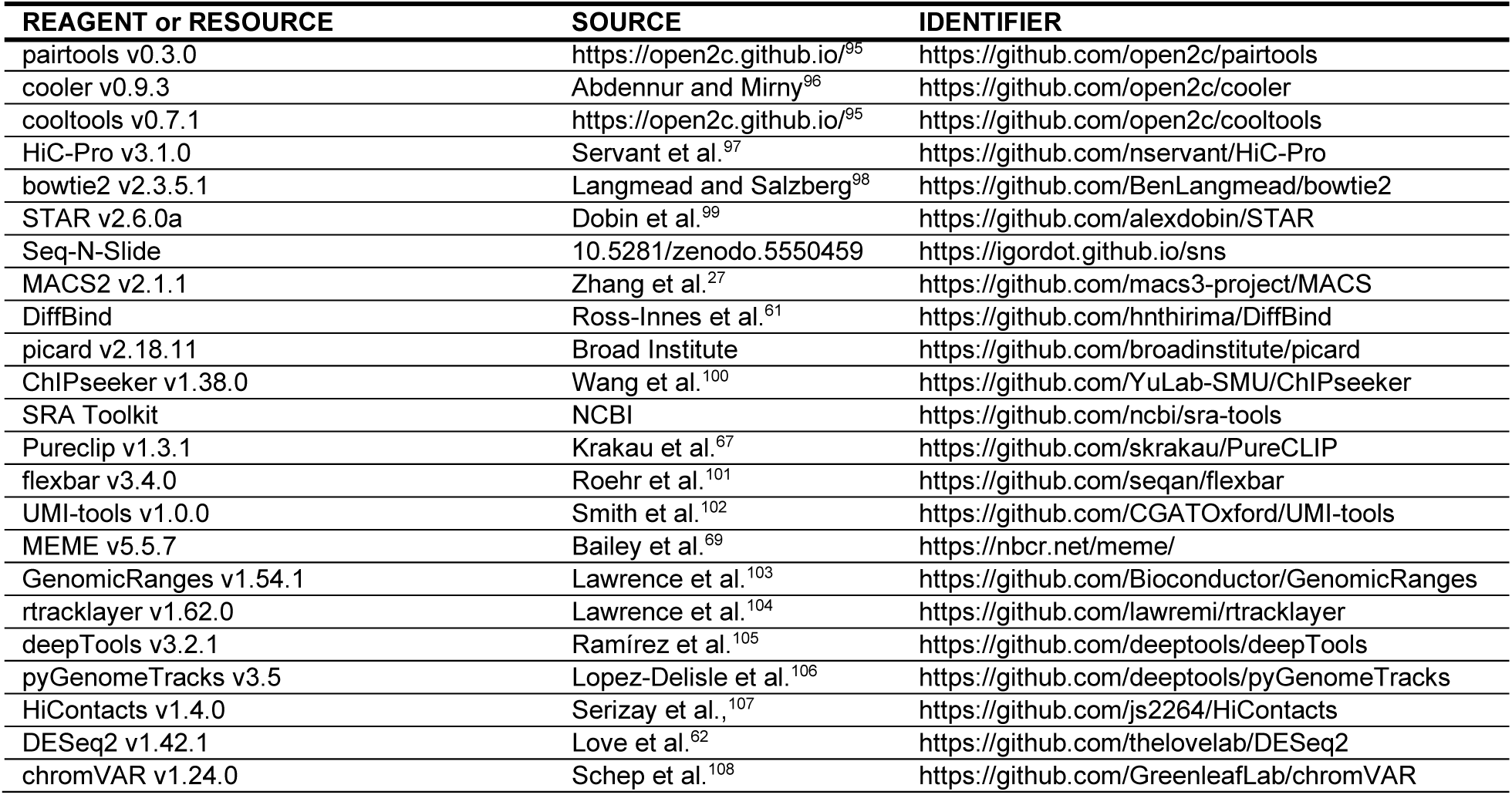

## EXPERIMENTAL MODEL DETAILS

The detailed cell lines are reported in key resources table. This study has been performed under compliance with ethical regulations and approved by NYU/NYULMC Institutional Biosafety Committee.

## METHOD DETAILS

### Cell Culture and NPC differentiation

E14Tga2 (ATCC, CRL-1821) mouse ESCs (E14) were cultured in N2B27 media [Advanced Dulbecco’s modified eagle medium (DMEM)/F12:Neurobasal-A (1:1) medium (ThermoFisher Scientific 12634028 and 10888022, respectively), with 3% ESC-grade fetal bovine serum (v/v, Gemini 900-108), 1x N2 (ThermoFisher Scientific 17502-048), 1x B27 (Thermo Fisher 17504-044), 2 mM L-glutamine (Sigma G7513), 0.1 mg/mL Penicillin-Streptomycin (Sigma P0781), 0.1 mM β-mercaptoethanol (ThermoFisher Scientific 21985023)] supplemented with leukemia inhibitory factor (homemade), 3 μM CHIR99021 (Tocris 4423) and 1 μM PD0325901 (Sigma PZ0162). Cells were grown on 0.1% gelatin-coated (Millipore ES-006-B) plates at 37°C and 5% CO2. All cell lines were routinely checked for mycoplasma contamination and in all cases were negative.

*In vitro* differentiation of ESCs to NPCs has been described previously^51^. Briefly, plates were coated overnight at 37°C with 10 ug/mL Laminin (R&D Systems 3446-005-01) in 1x Phosphate Buffered Saline (PBS). ESCs were dissociated with TrypLE Express (Gibco 12605-028). Cells were seeded on laminin-coated plates at a pre-titrated density of ∼15,000/cm^2^. Titration of seeded ESCs is highly recommended due to variation between labs and cell lines. Cells were differentiated for 2 days in N2B27 media. NPC differentiation was confirmed by immunostaining with Sox1 antibody (R&D Systems AF3369).

For CTCF rescue experiments, E14 CTCF-AID2 degron; dox inducible CTCF-WT or ZF1 mutant rescue cells were treated with 1 uM 5-Ph-IAA (Bio Academia 30-003) and 1 ug/mL doxycycline (Sigma D5207) for 24 hours and then harvested for downstream applications.

### CRISPR-Cas9 genome editing

Single-guide RNAs (sgRNAs) were designed using CRISPR design tools in Benchling (https://www.benchling.com). sgRNA oligos (see **Table S9**) were cloned into SpCas9-expressing PX330, PX458, or PX459 vectors (Addgene 42230, 48138, 62988, respectively) as previously described (https://www.addgene.org/crispr/zhang/). Briefly, vectors were digested with BbsI-HF (NEB R3539). Forward and reverse sgRNA oligos were annealed together and phosphorylated using T4 Polynucleotide Kinase (PNK) (NEB M0201). A 1:100 dilution of the resulting phosphorylated oligo duplex was ligated to the BbsI-digested vector using T4 DNA ligase (NEB M0202). Donor plasmids were generated as described in the “Generation of plasmids and cell lines” section below (see also **Table S9** for cloning details). Plasmids were transformed into XL10 Gold (Agilent 50-125-094) or Stellar Competent Cells (Takara Bio 636766) at 1:10 dilution. All plasmids were confirmed by Sanger sequencing, which was done by Azenta Life Sciences or Psomagen. The sequencing results were aligned to the reference DNA using SnapGene.

For all transfections, ESCs at 70-80% confluency in 6-well plates containing 1 mL of media were used. Cells were transfected with 1-2 µg of donor plasmid and 0.5 µg of sgRNA plasmid, using 6 µL of Lipofectamine® 2000 (ThermoFisher Scientific 11668027) in 100-150 µL of Opti-MEM® Medium (ThermoFisher Scientific 31985062). Cells were incubated in transfection mix for 1-2 days prior to undergoing antibiotic or fluorescent selection. Cells were then FACS-sorted into 96-well plates (1 cell per well) using BD FACSAria II cell sorter. Single-cell clones were expanded and genomic DNA (gDNA) was obtained using DirectPCR Lysis Reagent (Viagen Biotech 102-T). Successful gene editing was confirmed by genotyping PCR (see **Table S9** for genotyping primers), and in addition, western blots or flow cytometry analysis when applicable. Mutant PCR products from genotyping were also confirmed by Sanger sequencing.

### Generation of plasmids and cell lines

#### Endogenous CTCF-ΔZF1 mutants

E14 ESCs were transfected with PX458-eBFP2-sgRNA_CTCF_ZF1 targeting CTCF ZF1 and single-stranded donor DNA (IDT). The homozygous mutant clones were confirmed by PCR and Sanger sequencing.

#### CTCF-AID2 degron and rescue lines

First, the endogenous CTCF locus was tagged with mAID-eGFP upstream of the N-terminus. The KL094 - PBS-mAID-EGFP CTCF donor plasmid was generated by Gibson cloning^109^ of a gBlock (mAID-eGFP flanked by CTCF homology arms, IDT) into the pBlueScript II SK (+) vector (see **Table S9** for details). E14 ESCs were transfected with donor plasmid and PX330-sgRNA006_CTCF sgRNA plasmid. eGFP-positive cells were sorted. Homozygous mAID-eGFP-CTCF ESCs (clone C7) were confirmed by genotyping and Western blot analysis.

Next, clone C7 was modified by knocking in mutant OsTIR(F74G) into the constitutively active Rosa26 locus. A plasmid containing WT OsTIR1 flanked by Rosa26 homology arms (Addgene 86233) was modified to express OsTIR1(F74G) (see **Table S9** for details). Clone C7 was transfected with the OsTIR1(F74G) donor plasmid and sgRNA plasmid targeting the Rosa26 locus (Addgene 64216). Cells were selected using a neomycin-resistance gene on the donor plasmid. Heterozygous OsTIR(F74G) (clone C7-E10) was confirmed by genotyping.

Although not used in this study, an mCherry reporter was knocked-in at the Hb9/Mnx1 locus (Hb9-T2A-mCherry) of the C7-E10 clone. Hb9 is a motor neuron-specific marker. Initially, a motor neuron differentiation system was intended for this study, but we switched to the NPC differentiation system due to its higher efficiency. Cells were transfected with double-stranded donor DNA (T2A-mCherry flanked by Hb9 homology arms) and PX458-BFP-sgRNA011_Hb9 sgRNA plasmid. Homozygous Hb9-T2A-mCherry was confirmed by genotyping. The Hb9 mCherry reporter was further verified by anti-Hb9 immunostaining and flow cytometry analysis, before and after motor neuron differentiation.

Next, clone C7-E10 was modified by knocking in a dox-inducible, Flag-Halo-tagged WT or ΔZF1 CTCF at the constitutively active Tigre locus. The WT and ΔZF1 donor plasmids were generated by modifying an existing plasmid (Addgene 156432) that contains a constitutively expressed rtta3G cassette and a dox-inducible CTCF, flanked by Tigre homology arms (see **Table S9** for details). The donor plasmid and sgRNA plasmid targeting the Tigre locus (Addgene 92144) were transfected into the C7-E10 clone. Cells with the dox-inducible knock-in were selected using a puromycin-resistance gene on the donor plasmid. Homozygous knock-in clones (clone C7-E10-D11 for WT and C7-E10-F6 for ΔZF1) were confirmed by genotyping. After selection, neomycin and puromycin resistance cassettes were excised using a plasmid containing FLP recombinase (Addgene 13793).

#### Podxl and Grb10 pA lines

The donor plasmids were generated by first cloning ∼1000 bp of *Podxl* or *Grb10* gDNA from E14 lysates into the pBlueScript II SK (+) vector (Agilent 212207) (see **Table S9** for details). Using Gibson cloning^109^, T2A-eGFP-SV40pA was then inserted in the middle of the *Podxl* or *Grb10* gDNA such that it was flanked by ∼500 bp of homology arms (see **Table S9** for details). E14 ESCs were transfected with donor plasmids and sgRNA plasmids targeting the corresponding gene (PX459-sgRNA047_Podxl or PX459-sgRNA048_Grb10). Cells were selected using a puromycin-resistance gene on the sgRNA plasmid. eGFP-positive cells were then sorted. The homozygous mutant clones were confirmed by PCR and Sanger sequencing.

### Immunostaining and flow cytometry

Cells were dissociated into single cells with TrypLE Express, washed with 1x PBS, and fixed with 4% paraformaldehyde for 10 minutes at r.t. Cells were washed with FACS buffer (4% Fetal Bovine Serum and 2 mM EDTA in 1x PBS) twice and permeabilized in 0.5% saponin in FACS buffer for 10 min at r.t. 0.5% saponin in FACS buffer was used for antibody incubation. Cells were incubated with Goat anti-Sox1 primary antibody (R&D Systems AF3369) overnight at 4°C (1:200 dilution from 500 ug/mL antibody stock). After washing, cells were incubated with anti-Goat IgG-AF647 secondary antibody (Invitrogen A32849) (1:1000 dilution) for 1 hr at 4°C.

For HaloTag labeling, cells were incubated in 37°C with 10 nM HaloTag-JF646 (Promega GA1120) in media for 30 min. Cells were washed 3 times with 1x PBS and further incubated in media at 37°C for 30 min. Cells were then dissociated with TrypLE express and analyzed by flow cytometry.

Immunostained and HaloTag-labeled cells were assayed with BD LSRII or Bio-Rad ZE5 flow cytometer and results were analyzed using FlowJo software.

### Whole cell extract and western blotting

A confluent 6 well plate was lysed with 200 uL cold RIPA buffer (20 mM Tris, pH 7.5, 150 mM NaCl, 1% SDS, 5% sodium deoxycholate, and 1% NP-40) supplemented with 1 ug/mL Aprotinin, Pepstatin A, Leupeptin, and 0.2 mM PMSF protease inhibitors (PI). The cell lysate was briefly probe-sonicated (40% amplitude, 5 strokes) and centrifuged at 15,000 g at 4°C for 10 min. The supernatant was collected, and protein concentrations were quantified via bicinchonic acid (BCA) assay (ThermoFisher Scientific A55861). Proteins were denatured in 1x Laemmli SDS-PAGE buffer (63 mM Tris-HCl, pH 6.8, 10% glycerol, 2% SDS, 0.0005% Bromophenol blue, and 0.1% β-mercaptoethanol) at 95°C for 5 min. 20 ug total proteins were separated using a 6%–15% SDS-PAGE gel and transferred onto a PVDF membrane. Membranes were blocked with 5% milk in TBST at r.t. for 30 min and incubated with primary antibody overnight at 4°C (see **Table S10** for list of antibodies). Membranes were washed 3 times with TBST and then incubated with horseradish peroxidase (HRP)-conjugated secondary antibodies (see **Table S10** for list of antibodies) for 1 hr at r.t. HRP enzyme activity was detected by enhanced chemiluminescence (ThermoFisher Scientific 32106) and exposure to film, or by chemiluminescence detection using ChemiDoc^TM^ MP imaging system (Bio-Rad).

### Micro-C

Micro-C was done as described previously for mammalian cells^54^. Briefly, cells in a confluent 15 cm tissue culture plate (∼30 million (M) cells) were dissociated with TrypLE Express and crosslinked with freshly made 3 mM ethylene glycol bis (EGS) in PBS (5 mL per 5M cells), incubated at r.t. for 45 min. Crosslinking reaction was quenched by adding Tris pH 7.5 to final 0.75 M concentration and incubated at r.t. for 5 min. EGS-crosslinked cells were washed twice with 0.5% BSA in PBS and subjected to the second crosslinking with 1% formaldehyde in PBS (5 mL per 5M cells), incubated at r.t. for 10 min. Crosslinking reaction was quenched again by adding Tris pH 7.5 to final 0.75 M concentration, incubated at r.t. for 5 min. Cells were washed twice with 0.5% BSA in PBS and aliquoted into 5M cells per 1.5 mL tube. Cell pellets were spun down, snap-frozen, and stored at −80°C.

Crosslinked cells (5M in 1.5 mL tubes) were thawed on ice and permeabilized in cold Micro-C Buffer (MB) #1 (50 mM NaCl, 10 mM Tris, pH 7.5, 5 mM MgCl_2_, 1 mM CaCl_2_, 0.2% NP-40, and 1x Protease Inhibitor Cocktail) for 20 min on ice. Chromatin from permeabilized cells was digested by adding micrococcal nuclease (MNase) (NEB M0247SVIAL) at 1:100 dilution and incubated at 37°C for 10 min, shaking at 850 rpm. Dilution of MNase was pre-titrated such that chromatin is digested efficiently into nucleosome-sized fragments. Titration is recommended due to enzyme lot variability. Enzyme activity was stopped by adding EGTA to a final concentration of 4 mM and incubated at 65°C for 10 min. MNase-digested chromatin was washed twice with ice-cold MB #2 (50 mM NaCl, 10 mM Tris, pH 7.5, and 10 mM MgCl2).

Digested chromatin fragments were then subjected to 3’-dephosphorylation and 5’-phosphorylation using 0.5 U/uL T4 PNK in 1x NEBuffer 2.1 (NEB B6002S) with 2 mM ATP and 5 mM DTT, incubated for 15 min at 37°C with interval mixing. 3’-end-chewing was done by adding 0.5 U/uL DNA Polymerase I Klenow Fragment and incubating another 15 min in 37°C. Blunt-end reaction was triggered by adding biotin-dATP, biotin-dCTP, dGTP, and dTTP to a final concentration of 66 uM and incubated at 25°C for 45 min. The reaction was stopped by adding 30 mM EDTA final concentration and incubating at 65°C for 20 min. Chromatin was washed once with cold MB #3 (50 mM Tris, pH 7.5, and 10 mM MgCl2).

Chromatin fragments with biotin-dNTPs were then ligated using 20 U/uL T4 DNA ligase (NEB M0202M), incubated at r.t. for at least 2.5 hr with slow rotation. Unligated ends containing biotin-dNTPs were then removed by 5 U/uL exonuclease III in 1x NEBuffer 1 (NEB B7001S), incubated at 37°C for 15 min. Chromatin was subjected to reverse crosslinking with 1 mg/mL Proteinase K (Sigma 3115879001), 1% SDS, and 100 ug/mL RNase A (ThermoFisher Scientific EN0531), incubated at 65°C overnight.

DNA was extracted by adding 1x volume Phenol:Chloroform:Isoamyl Alcohol (25:24:1) solution (Thermofisher Scientific 15593049) and isolating aqueous phase after centrifugation. DNA was purified from the aqueous phase using a PCR purification kit (Zymo Research DCC-5 D4013) following manufacturer’s instructions. Separation of monomers and dimers was done by running purified DNA on 3% TBE agarose gel. 200-400 bp DNA was then extracted from the gel by Zymo Gel DNA recovery kit (Zymo Research D4008).

DNA with biotin-dNTPs was captured by Dynabeads® MyOne Streptavidin C1 (ThermoFisher Scientific 65001). Standard library preparation protocol including end-repair, A-tailing, and adaptor ligation was performed on beads with enzymes from NEBNext Ultra II DNA Library prep kit for Illumina (E7645S). NEBNext Multiplex Oligos for Illumina^®^ (Dual Index Primers Set 1, E7600S) were used for barcoding. Sequencing library was amplified by Kapa HiFi PCR enzyme (KAPA KK2601). Libraries were verified by High Sensitivity D1000 ScreenTape (Agilent 5067-5584) using the 4200 TapeStation System (Agilent). Libraries were sequenced as 2 x 50 or 2 x 150 bp paired-end reads on the Illumina Novaseq 6000 platform or Novaseq X platform, ensuring sufficient read depth of combined replicates for analysis (see **Table S8** for the number of reads in each sample replicate).

### ChIP-seq

ChIP-seq experiments were performed as described previously^110^. Briefly, cells in a confluent 15 cm tissue culture plate were washed once with PBS and fixed for 10 min with 1% formaldehyde in 15 mL fixation buffer (10 mM HEPES, pH 7.5, 15 mM NaCl, 0.15 mM EDTA, and 0.075 mM EGTA in DMEM). Formaldehyde was quenched with 125 mM glycine, incubated for 5 min at r.t. Cells were washed and scraped in PBS and transferred to 15 mL tubes. Nuclei were isolated using buffers supplemented with 1x protease inhibitor cocktail (Roche 11873580001) in the following order: 5 mL LB1 (20 mM Tris, pH 7.5, 10 mM NaCl, 1 mM EDTA, and 0.2% NP-40; 10 min on ice), 5 mL LB2 (20 mM Tris, pH 7.5, 200 mM NaCl, 1 mM EDTA, and 0.5 mM EGTA; 10 min on ice), and 1.2 mL LB3 (20 mM Tris, pH 7.5, 150 mM NaCl, 1 mM EDTA, 0.5 mM EGTA, 1% Triton X-100, 0.1% sodium deoxycholate, and 0.1% SDS). Nuclei in LB3 were transferred to 15 mL Bioruptor Pico tubes (Diagenode C30010017) containing 800 mg sonication beads. Chromatin was fragmented in LB3 to an average size of 200-250 bp using a Bioruptor Pico (Diagenode). Insoluble material was pelleted by centrifuging at 15,000 g for 10 minutes at 4°C, and supernatant was kept. Supernatant containing sonicated chromatin were pre-cleared with 25 uL Protein G Dynabeads (Invitrogen 10004D), incubated rotating at 4°C for 1 hr. For each reaction, 2-4 ug antibody (see **Table S10** for list of antibodies) was incubated with 10 ul Protein G Dynabeads in LB3, rotating at 4°C for 3 hr. Antibody-bead complexes were added to 300 ug of chromatin and incubated rotating at 4°C overnight. 30 ug chromatin was used as input control. Beads were then washed 4 times with RIPA buffer (50 mM HEPES, pH 7.5, 500 mM LiCl, 1 mM EDTA, 0.5% sodium deoxycholate, and 1% NP-40) and washed once with TE50 buffer (10 mM Tris, pH 8.0, 1 mM EDTA, and 50 mM NaCl). The beads-bound chromatin was eluted in 125 uL elution buffer (50 mM Tris, pH 8.0, 10 mM EDTA, and 1% SDS), incubated at 65°C for 1 hr. Eluted chromatin was de-crosslinked using 0.6 mg/mL proteinase K and 0.1 mg/mL RNase A, incubated overnight at 55°C, shaking 1000 rpm. 1x volume Phenol:Chloroform:Isoamyl Alcohol (25:24:1) solution was added to the elution and aqueous phase was isolated after centrifugation. DNA was purified from the aqueous phase using a PCR purification kit (Zymo Research DCC-5 D4013) following manufacturer’s instructions. Libraries were prepared using NEBNext Ultra II DNA Library prep kit for Illumina (E7645S) following manufacturer’s instructions but using ½ volume for all the reactions. NEBNext Multiplex Oligos for Illumina^®^ (Dual Index Primers Set 1, E7600S) were used for barcoding. Fragments of 200-700 bp were size-selected using Agencourt AMPure XP beads. Libraries were quantified with High Sensitivity D1000 ScreenTape and TapeStation System. Libraries were sequenced as 2 x 50 bp or 2 x 150 bp paired-end reads on the Illumina Novaseq X platform.

### ChIP-MS

Cells were trypsinized from tissue culture plates, washed with 1x PBS and pelleted. Nuclei were extracted by adding HMSD buffer (20 mM HEPES, pH 7.5 at 4°C, 5 mM MgCl2, 85.5 g/L sucrose, 25 mM NaCl, and 1 mM DTT) supplemented with protease inhibitors (PI) (0.2 mM PMSF, 1 mg/mL Pepstatin A, 1 mg/mL Leupeptin, and 1 mg/mL Aprotinin), and rotated in a cold room for 30 min. Nuclei were spun down. The resulting pellet was resuspended in Buffer A (10 mM Tris, pH 8.0, 1.5 mM MgCl_2_, 10 mM KCl, and 0.2% NP-40) supplemented with PI. Nuclei were rotated in Buffer A in cold room for 30 min. The nuclear pellet was spun down. 1 mL benzonase buffer (20 mM Tris, pH 8.0, 100 mM NaCl, and 2 mM MgCl_2_) with 1 uL Benzonase (5U/uL stock) (Sigma E8263) was added to chromatin pellet and incubated in cold room rotating to digest overnight. Insoluble material was pelleted, and supernatant was kept for anti-Flag immunoprecipitation of Flag-halo tagged CTCF rescues. Total protein was quantified with BCA assay and 400-600 ug of proteins were used for immunoprecipitation for each sample replicate. 10 uL of anti-flag agarose beads (Sigma A2220) was added to supernatant and incubated rotating in cold room overnight. After washing the beads 3 times with benzonase buffer, Flag-halo-CTCF was eluted with 10 uL 1x Flag peptide (5 mg/mL stock) (Sigma F3290) in 90 uL BC100 buffer (40 mM Tris, pH 7.3, 100 mM NaCl, 5 mM MgCl_2_, and 5% glycerol) supplemented with PI. A first elution was done overnight at 4°C, and a second elution done for 2-3 hr at 4°C. Both elutions were combined. 10-20 uL of pooled elution was separated by SDS– PAGE, using 6%–15% SDS-PAGE gradient gels. Silver staining and western blots were done to confirm the presence of the bait (CTCF), as well as a known protein interactor (Rad21). Protein elutions were sent to Rutgers University Mass Spectrometry Facility for standard LC-MS/MS procedure. The facility performed differential spectral count analysis as previously described^111^, and results are reported in **Table S1, S2**.

### Bulk RNA-seq

Cells from a confluent 6-well plate were dissociated with TrypLE Express and washed once with 1x PBS. RNA was purified using RNeasy Plus Mini Kit (Qiagen 74136). RNA integrity number >7 was verified using Agilent RNA ScreenTape (Agilent 5067-5579). rRNA was depleted from 1 ug starting total RNA using NEBNext rRNA Depletion Kit v2 (NEB E7405) following manufacturer’s instructions. cDNA was synthesized using NEBNext Ultra II RNA first strand synthesis module (NEB E7771) (with random hexamers) and NEBNext Ultra II Directional RNA second strand synthesis module (NEB E7550) following manufacturer’s instructions. Libraries were prepared using NEBNext Ultra II DNA Library prep kit for Illumina (E7645S) in conjunction with NEBNext Multiplex Oligos for Illumina (E7600S or E7335S) for barcoding. Libraries were purified with Agencourt AMPure XP beads. Libraries were quantified with High Sensitivity D1000 ScreenTape and TapeStation System. Libraries were sequenced as 2 x 50 bp paired-end reads on the Illumina Novaseq 6000 platform or Novaseq X platform.

### RT-qPCR

Cells from a confluent 6-well plate were dissociated with TrypLE Express and washed once with 1x PBS. RNA was purified using RNeasy Plus Mini Kit (Qiagen 74136). cDNA synthesis was done on 2 ug RNA using Superscript III Reverse Transcriptase (Invitrogen 18080044) and Oligo(dT)_20_ primer (ThermoFisher Scientific 18418020) in 20 uL total volume. cDNA was diluted with 380 uL nuclease-free water. 4 uL of diluted cDNA was used for qPCR. RT-qPCRs were performed in triplicates using PowerUp SYBR Green Master Mix (ThermoFisher Scientific A25742) on a CFX384 Touch Real-Time PCR detection system (Bio-Rad). GAPDH-normalized relative expression and statistical analyses were calculated using CFX Maestro Software 2.3. The primers used are listed in **Table S9**.

### CLIP-seq

CLIP-seq was based on a previously described iCLIP2 protocol^66^ with modifications. A confluent 15 cm tissue culture plate was washed once with 1x PBS. 10 mL 1x PBS was added to the plate and cells were UV-crosslinked on plate with opened lid, at 400 mJ/cm^2^ using SpectroLinker XL-1000 (254 nm). Two 15 cm plates were scraped and combined. Cells were pelleted, snap-frozen and stored at-80°C. ±UV cell pellets were resuspended in 20 mL HMSD (20 mM HEPES, pH 7.5, 5 mM MgCl_2_, 25 mM NaCl, 85.5 g/L sucrose, and 1 mM DTT) supplemented with protease inhibitors (0.2 mM PMSF, 1 mg/mL Pepstatin A, 1 mg/mL Leupeptin, and 1 mg/mL Aprotinin), and rotated in cold room for 25 min. After washing with 1x PBS, nuclei were resuspended in 1 mL cold PBS-lysis buffer (1 mM MgCl_2_, 0.1 mM CaCl_2_, 0.5% sodium deoxycholate, and 0.5% NP-40 in 1x PBS) supplemented with 1x EDTA-free protease inhibitor cocktail (PIC) (Sigma 11873580001), 40 U/mL Protector RNase inhibitor (Sigma 3335402001), 1 mM DTT, and rotated in cold room for 25 min. 20 uL of Turbo DNase (ThermoFisher Scientific AM2238) was added and lysates were incubated at 37°C for 30 min, shaking 1100 rpm. Partial RNA digestion was done with a pre-titrated volume of dilute RNase I (ThermoFisher Scientific AM2295), incubated for 3 min at 37°C and immediately transferred to ice. 2 uL of SUPERaseIN (ThermoFisher Scientific AM2696) and 0.1% final concentration of SDS was added to stop RNA digestion. Lysates were spun down and supernatant was transferred to new 1.5 mL tubes. Flag-tagged CTCF from supernatant was immunoprecipitated with 5 uL anti-Flag magnetic beads (Millipore Sigma M8823), rotating in cold room for 2 hr. After 4 washes with PBS high-salt wash buffer (1 M NaCl, 0.1% SDS, 0.5% sodium deoxycholate, and 1% NP-40 in 1x PBS) and then 4 washes with PBS low-salt wash buffer (150 mM NaCl, 0.1% SDS, 0.5% sodium deoxycholate, and 1% NP-40 in 1x PBS), beads were treated with 0.1 U/uL Turbo DNase with 1x protease inhibitor cocktail, 0.4 U/uL Roche Protector RNase inhibitor, and 0.1 U/uL SUPERaseIn RNase inhibitor in 1x DNase buffer (ThermoFisher Scientific AM 2239). Beads were incubated at 37°C for 30 min, shaking at 850 rpm. Subsequently, RNAs were either radiolabeled for visualization or purified for cDNA library preparation.

For radiolabeling, on-beads RNA was 5’-labeled with 0.45 mCi/mL ATP, [g-^32^P] (Perkin Elmer BLU002Z250UC) using T4 PNK (NEB M0201L) in the supplied PNK buffer, supplemented with 0.9 U/uL Roche Protector RNase inhibitor. Standard precautions and proper waste disposal were observed when working with radioactive ATP. CTCF was eluted from beads twice using 1x flag peptide (5 mg/mL stock) (Sigma F3290) diluted 1:10 in PK buffer (100 mM Tris, pH 8.0, 150 mM NaCl, 5 mM EDTA, and 0.1% SDS). Proteins were denatured with 1x NuPAGE™ LDS sample buffer (ThermoFisher Scientific NP0007), incubated at 95°C for 5 min. The radiolabeled protein-RNA complexes were separated by electrophoresis using a 6% Bis-Tris gel and transferred to a nitrocellulose membrane. The radioactive membrane was exposed to film overnight at-80°C. CTCF western blots were done on the same membrane as described in the “Whole cell extract and western blotting” section above.

For cDNA preparation, on-beads RNA was dephosphorylated at the 3’ end using T4 PNK and ligated to a DNA oligo (L3-App) adapter (see **Table S9**) using T4 RNA Ligase 1 (NEB M0204L). CTCF was eluted from beads twice using 1x flag peptide (5 mg/mL stock) (Sigma F3290) diluted 1:10 in PK buffer (100 mM Tris, pH 8.0, 150 mM NaCl, 5 mM EDTA, and 0.1% SDS). Elutions were pooled and proteins were denatured and digested with 4 mg/mL proteinase K incubated in 55°C for 20 min, shaking 1200 rpm. 7M final concentration of urea was added and incubated at 55°C for another 20 min. RNA-DNA hybrids were extracted using phenol:chloroform:isoamyl alcohol, separating the aqueous phase using phase lock gel tubes (VWR 10847-802). From the aqueous phase, RNA-DNA hybrids were purified by ethanol precipitation overnight at-20°C. RNA-DNA pellet was washed with ethanol and resuspended in nuclease-free water. Reverse transcription (RT) was done using Superscript III Reverse Transcriptase (Invitrogen 18080044). The primer used for RT (RToligo) (see **Table S9**) was complementary to the DNA oligo (L3-App) previously ligated to RNAs. Despite extensive digestion by proteinase K of crosslinked protein-RNA complexes, a small polypeptide remains covalently linked to RNA, which prematurely terminates RT, thereby providing positional information of the crosslinked nucleotides in the resulting cDNA library. Synthesized cDNAs were extracted using Dynabeads MyONE Silane beads (ThermoFisher Scientific 37002D). A second adapter ligation, with barcodes (see **Table S9**), was done on the 3’ end of cDNA using T4 RNA Ligase 1. Adapter-ligated cDNAs were extracted using Dynabeads MyONE Silane beads. The adapter ligations are designed in such a way that sequencing reads start at the position where the cDNA was truncated during RT^66^. Adapters contained unique molecular identifiers (UMIs) for duplicate removal during analysis. A cDNA pre-amplification was done on beads for 6 cycles using 2x KAPA HiFI Hot Start Mix (KAPA KK2601). The primers used were P5Solexa_s and P3Solexa_s (see **Table S9**). ProNex® Size-Selective Purification System (Promega NG2001) was used to size select the pre-amplified cDNA and remove primer-dimers. A second PCR amplification was done for 9 cycles using primers P5Solexa and P3Solexa (see **Table S9**). Libraries were size selected (200-500 bp) with Agencourt AMPure XP beads. Libraries were quantified with High Sensitivity D1000 ScreenTape and TapeStation System. Libraries were sequenced as 100 bp single-end reads on the Illumina Novaseq 6000 platform.

## COMPUTATIONAL ANALYSIS

### Micro-C analysis

Micro-C analysis was performed according to the pipeline described by Dovetail Genomics in https://micro-c.readthedocs.io/en/latest/index.html. Briefly, multiple pairs of fastq files (2 biological replicates per condition) were aligned to the mm10 reference genome using BWA-MEM split alignment. Pairtools was used for the next steps: identifying valid ligation events (pairs), removing PCR duplicates, generating and down-sampling the final pairs files. Pairs files were down-sampled such that all samples being compared together had equal number of valid pairs. The down-sampled pairs files were used for downstream analyses. Cool and mcool files were generated using cooler^96^. Hic contact matrices were generated using Juicer Tools^57^. APA plots and scores were generated using Juicer Tools^57^. Micro-C contact heatmaps were generated using HiContacts^107^ and pyGenomeTracks^106^.

Compartments were characterized as described previously^112^. HiC-Pro^97^ was used to build contact matrices, perform iterative correction and eigenvector decomposition (ICE) normalization. HiC-Pro was used to generate ICE-normalized matrixes for 5-kb and 10-kb resolutions and annotation files that indicated the genomic bins. TADs were identified using Arrowhead^57^ at 10kb resolution with default options. GENOVA^58^ was used to visualize ATA. Insulation score bigwig files were generated from cool files using cooltools^113^. Deeptools^105^ was used to plot insulation score profiles at CTCF ChIP-seq peaks.

### Chromatin loop analysis

Chromatin loop analysis of Micro-C data was performed as previously described^114^. Fithic2^56^ was used to determine the contact counts of each pair-wise 5-kb bins, and calculate the likelihood of a contact to be significantly higher than expected for its genomic distance (*q*-value). Loops were classified between pair-wise samples (e.g. WT vs ΔZF1, ESC vs NPC) as “sample1”-specific, “sample2”-specific, or common. A loop was considered sample1-specific if it was found to be significant only in sample1 but not in sample2, when using a *q*-value cutoff of 0.01. To add more stringency, a loop was considered to be significantly different only if the log_2_ fold-change of the contact counts were ≤-1 or ≥ 1.

### RNA-seq analysis

RNA-seq fastq files were processed using the route “rna-star” and “rna-star-groups-dge” from the Slide-n-Seq (sns) pipeline: https://igordot.github.io/sns/. Processing steps include alignment to the mouse reference genome (mm10) using the STAR aligner with default parameters. Counts were obtained using featureCounts. Deeptools was used to generate bigwig files from the merged bam files of replicates for the same condition. Bigwig tracks were visualized using IGV and pyGenomeTracks. Downstream analysis, including normalization and differential expression analysis, was performed using DESeq2^62^. Genes were categorized as differentially expressed if log_2_ fold-change ≤-0.5 or ≥ 0.5, and adjusted p-value ≤ 0.05. Complete differential gene expression analysis is listed in **Table S3, S4**. GO term enrichment analysis of differentially-expressed genes was performed using PANTHER^63^. Complete GO Terms are listed in **Table S5**.

### ChIP-seq analysis

ChIP-seq analysis was done as previously described^110^. Reads were aligned to the mouse reference genome mm10, using Bowtie2 with default parameters. Reads of quality score less than 30 were removed using samtools and PCR duplicates were removed using picard. Regions in mm10 genome blacklist was removed using bedtools to generate the final BAM files. The final BAM files for replicates of same condition were merged using samtools. Bigwig files were generated from merged BAM files using deeptools with parameters:--binSize 50--normalizeUsing RPKM--ignoreDuplicates -- ignoreForNormalization chrX--extendReads 250. Bigwig files were visualized using IGV and pyGenomeTracks. Peaks were called using MACS2^60^ with parameters-g mm-B-f BAMPE--keep-dup all-q 0.01. Using MACS2-called peaks as input, differential ChIP-seq peaks and consensus peaks were identified using DiffBind^61^. A peak was considered significantly different between samples if log_2_ fold-change ≤-1 or ≥ 1, and adjusted p-value ≤ 0.05. Heatmaps were generated using the functions computeMatrix followed by plotHeatmap from deepTools. Violin plots for ChIP-seq signal were prepared using an in-house R code as follows: regions of interest (i.e., TSS-TES of genes) are imported from a BED-file as Granges object using GenomicRanges package. The number of ChIP-seq reads overlapping each region is counted from the BAM files using the getCounts function from the chromVAR package. Counts are Deseq2 normalized and the mean count for each region is calculated from all replicates of each sample. The resulting counts file was used as input to ggplot2 for Violin plots.

### CLIP-seq analysis

CLIP-seq analysis was performed using publicly available code that was described previously^68^. Briefly, FastQC was used to assess the quality of sequencing reads. Flexbar was employed to demultiplex reads based on barcodes and trim adapters. The reads were then aligned to the mm10 reference genome using STAR. PCR duplicates were removed using unique molecular identifiers (UMIs) with UMI-tools. As described in the “CLIP-seq” protocol (above), reads start at the position where the cDNA was truncated during reverse transcription, making the position upstream of the 5’ end of the read the ‘crosslinked nucleotide’. Bedtools was used to convert BAM files to BED files, shift the BED intervals by one base pair in the 5’ direction, and extract the 5’ positions of the new intervals. These positions were then piled up into BEDGraph files, separately for each strand. BEDGraph files were converted to BigWig files using bedGraphToBigWig. Bigwig files were visualized using IGV and plotted with pyGenomeTracks. For peak calling, BAM files from replicates of the same condition were merged and used as input for PureCLIP^67^, which outputs individual crosslink sites. PureCLIP-called crosslink sites were post-processed as described previously^68^, to obtain equal-sized CLIP peaks of 300 nt. For each called peak, the number of crosslink events per replicate was determined. A minimum cutoff (10th percentile of the number of crosslinked sites) was calculated for each replicate. A peak was considered reproducible if it met the cutoff in a sufficient number of replicates (at least 3 out of 4 replicates, except for NPC WT, where the criterion was 2 out of 3 replicates). Peak calling was also performed for non-crosslinked negative controls, and peaks with at least 20% overlap with negative control peaks were filtered out. Reproducible CLIP peaks were assigned to gene names and gene types based on GENCODE annotations (GRCm38.p4). *De novo* RNA motif analysis using the reproducible CLIP peaks as input was performed using MEME^69^. The assignment of CLIP peaks to genomic features (e.g., promoters, introns, exons) was done using ChIPseeker^100^. Differential CLIP genes between ESCs and NPCs were identified by combining unique annotated CLIP genes from both cell types and retrieving the TSS-TES regions as a BED file. The BED file was imported as a GRanges object using GenomicRanges package on R, and the number of CLIP-seq reads overlapping each region was counted from the BAM files using the getCounts function from the chromVAR package. DESeq2 was used to normalize the read counts and identify differential genes (log_2_ fold-change cutoff of ≤-1 or ≥ 1, adjusted p-value ≤ 0.05). Heatmaps of CLIP peaks and CLIP-seq signal from TSS-TES regions of annotated genes were generated from bigwig files using deepTools. Venn diagrams were drawn using VennDiagram package on R studio.

### Statistical Analysis

Statistical analysis related to experiments have been described above in each section.

## SUPPLEMENTARY FIGURES

**Figure S1.**
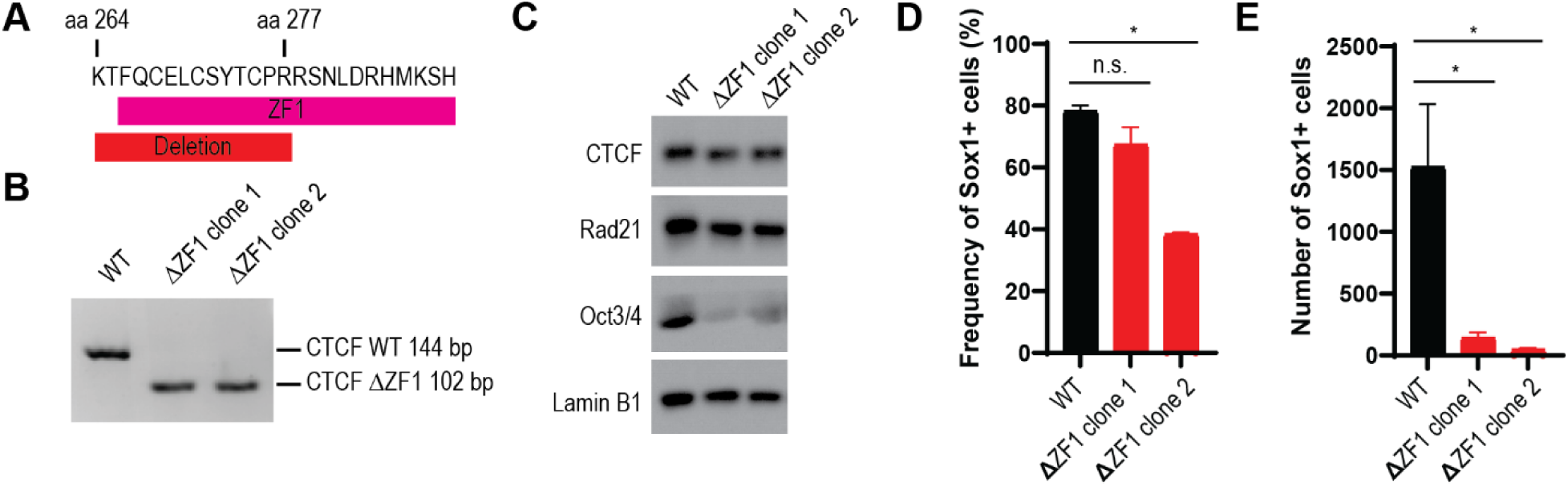
Endogenous CTCF-ΔZF1 mutants do not efficiently differentiate into NPCs, related to. Figure 1. (A) Schematic of 14-amino acid (aa) deletion of CTCF-ZF1 in ESCs. (B) Genotyping of endogenous CTCF-ΔZF1 clones. Deletion was confirmed by DNA sequencing of purified bands. (C) Western blot of parental WT ESCs and ΔZF1 clones. (D) Percentages of Sox1+ cells after 2 days of NPC differentiation. Cells were fixed and immunostained with anti-Sox1 antibody and analyzed by flow cytometry. Data are represented as mean ± SEM, p-values were determined using Dunnett’s multiple comparison test, n.s.=not significant, * p < 0.05, N=2. (E) Number of Sox1+ cells per 5,000 seeded ESCs, after 2 days of NPC differentiation. Cells were fixed and immunostained with anti-Sox1 antibody and analyzed by flow cytometry. Cell numbers were normalized by cell counting beads. Data are represented as mean ± SEM, p-values were determined using Dunnett’s multiple comparison test, * p < 0.05, N=2.

**Figure S2.**
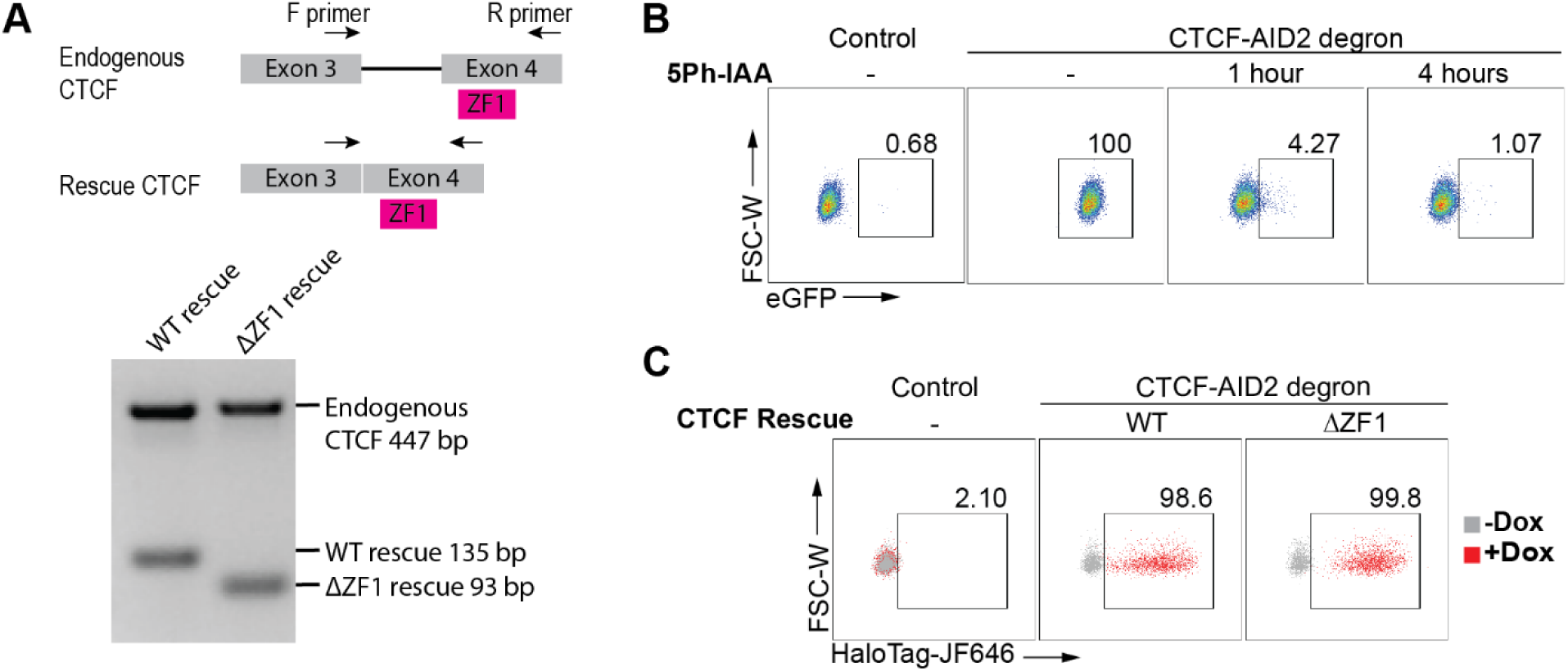
Validation of CTCF-AID2 and rescue lines, related to. Figure 1. (A) The schematic (above) shows the genotyping strategy with the arrows indicating forward (F) and reverse (R) primers. Primers amplify both endogenous and rescue CTCF but result in different DNA base pair (bp) lengths. Below is the genotyping of CTCF-AID2 and rescue lines. (B) CTCF-AID2 degron line after no treatment, 1 hr, or 4 hr of treatment with 5-Ph-IAA. eGFP fluorescence was then analyzed by flow cytometry. The negative control is the parental E14 WT cell line. Numbers are the percentages of cells within the boxed area among all cells in the plot. (C) CTCF-AID2 degron line after no treatment, or 24 hr of treatment with dox. Cells were labeled with HaloTag-JF646. JF646 fluorescence was then analyzed by flow cytometry. The negative control is the parental E14 WT cell line. Numbers are the percentages of cells within the boxed area among all dox-treated cells (in red) in the plot.

**Figure S3.**
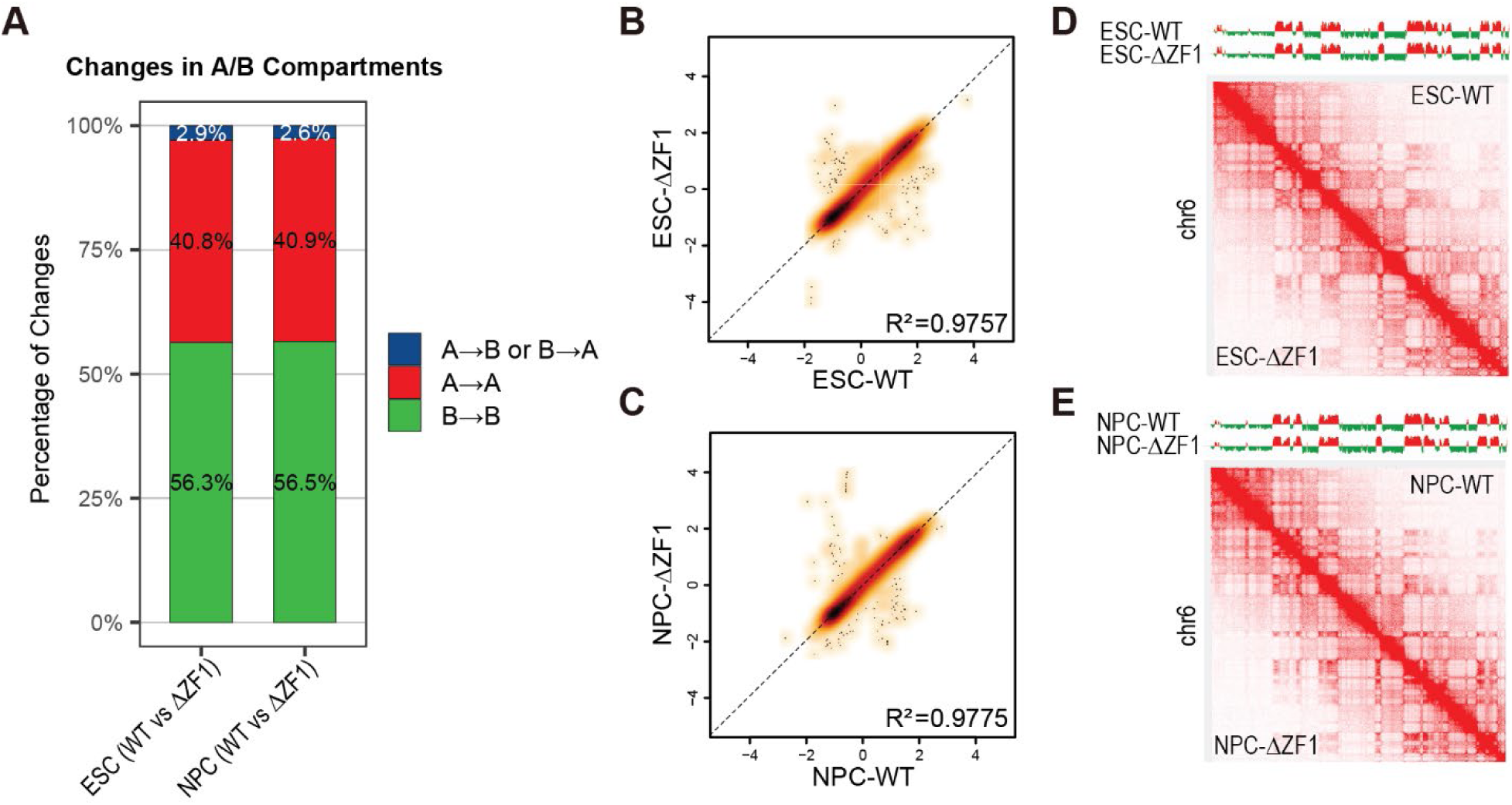
Genomic compartmentalization is minimally changed in ΔZF1 for both ESCs and NPCs, related to. Figure 2. (A) Bar graphs showing the percentage of compartment changes from WT to ΔZF1. (B-C) Scatter plots comparing the first eigenvector values (equivalent to first principal component) between WT and ΔZF1 in (B) ESCs and (C) NPCs. The correlation coefficient (R^2^) was calculated. Dashed lines represent linear regression lines. (D-E) The first eigenvector tracks (top) and Micro-C heatmaps (bottom) comparing WT and ΔZF1 in (D) ESCs and (E) NPCs.

**Figure S4.**
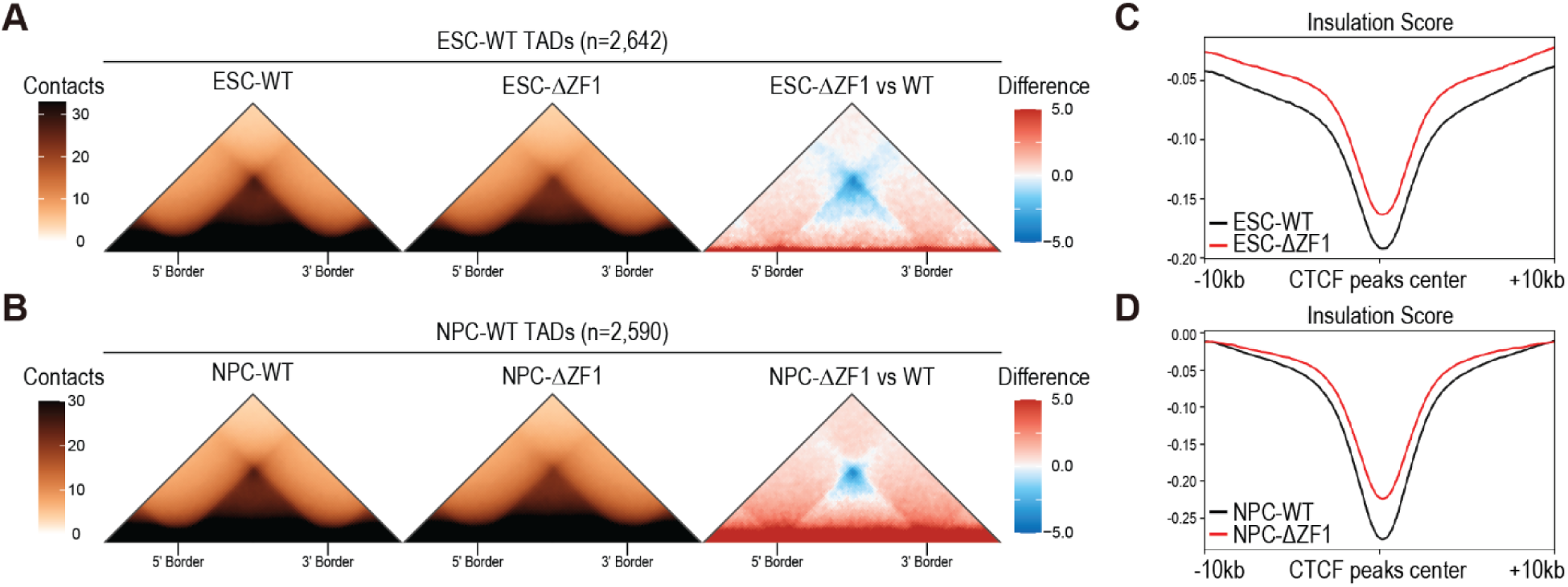
TADs and TAD boundaries are weakened in ΔZF1 for both ESCs and NPCs, related to. Figure 2. (A-B) Aggregate TAD analysis (ATA) of WT (left), ΔZF1 (middle), and the difference between ΔZF1 and WT (right) in (A) ESCs and (B) NPCs. (C-D) Diamond insulation scores (1kb resolution, 10kb window size) of WT and ΔZF1 centered at CTCF ChIP-seq peaks in (C) ESCs and (D) NPCs.

**Figure S5.**
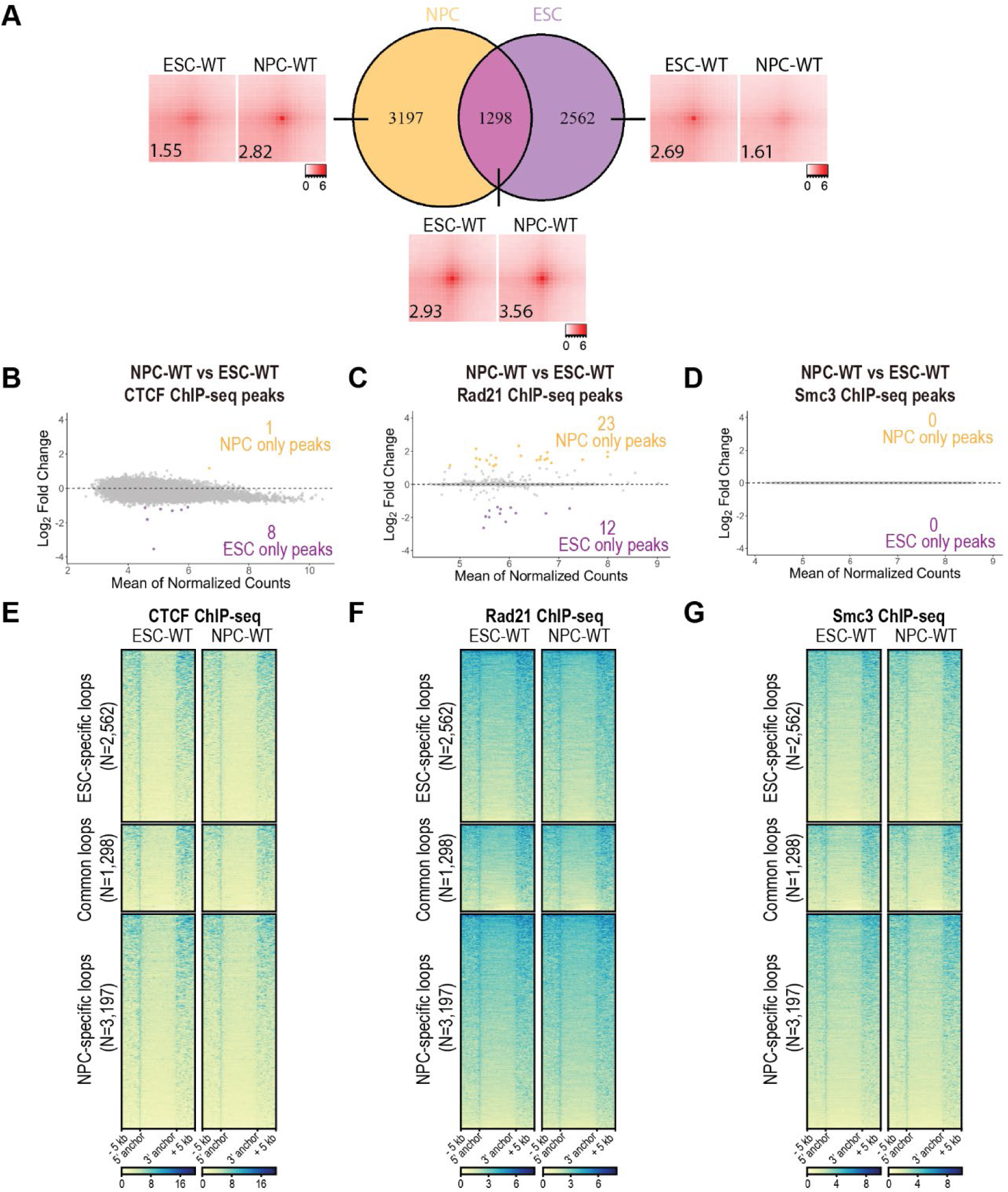
ESCs and NPCs exhibit differential chromatin loops, but CTCF and cohesin binding are mostly cell-type-invariant, related to. Figure 2. (A) Venn diagram of NPC and ESC CTCF anchors. APA plots show aggregated peaks from each of the NPC-specific, ESC/NPC common, and ESC-specific loop subsets. Numbers indicate APA scores. (B, C, D) DiffBind MA plot of differentially called (C) CTCF, (D) Rad21, and (E) Smc3 ChIP-seq peaks between NPC-WT vs ESC-WT. Adjusted p-value cutoff: ≤ 0.05, log_2_ fold-change cutoff: ≥ 1, ≤-1, CTCF N=4, Rad21 N=2, Smc3 N=2. (E, F, G) ChIP-seq heatmaps of (E) CTCF, (F) Rad21, and (G) Smc3, comparing chromatin binding in ESC-WT and NPC-WT. Each row is a loop anchor coordinate, and the heatmap is clustered based on whether the anchors are ESC-specific, ESC/NPC common, or NPC-specific.

**Figure S6.**
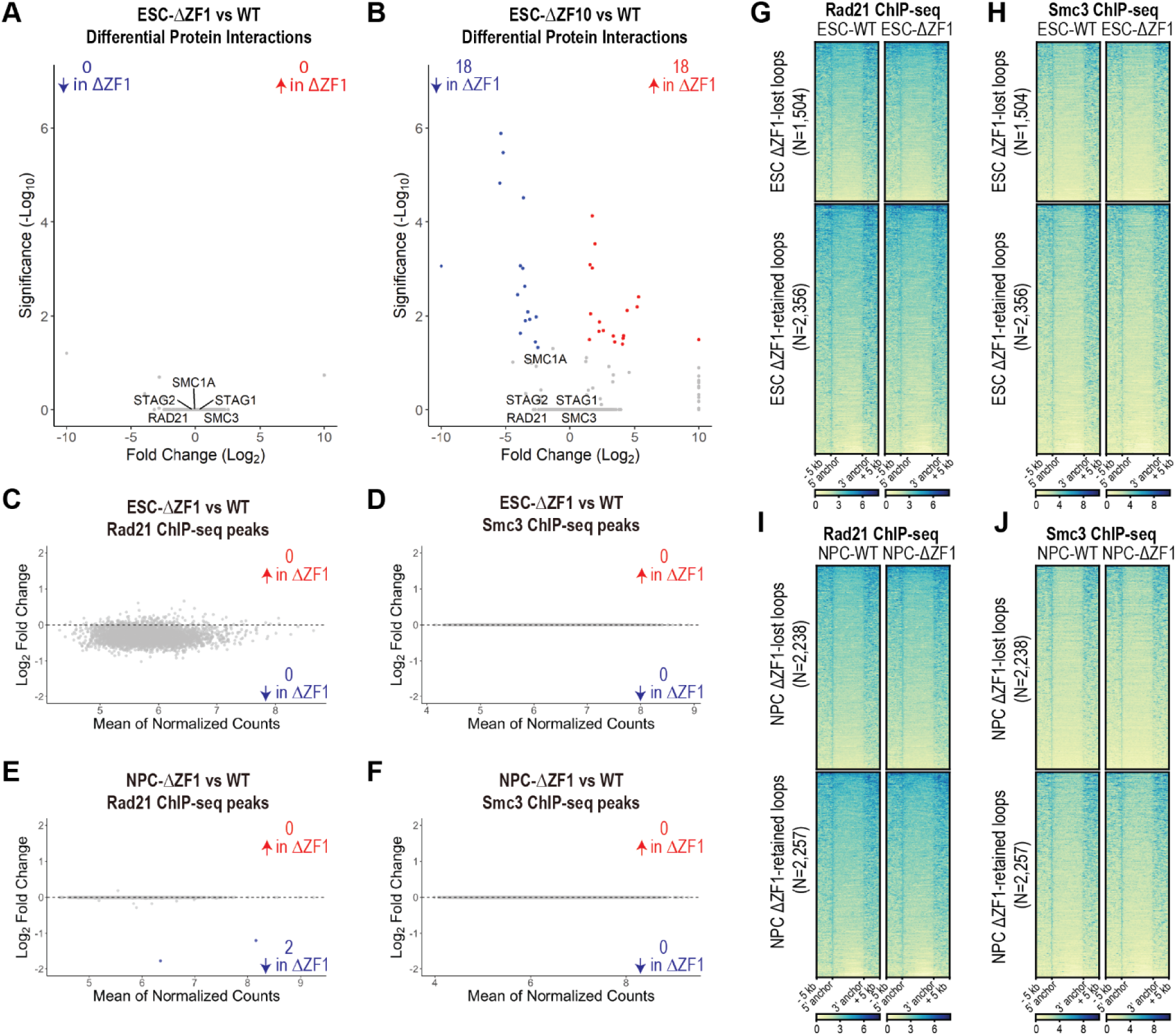
CTCF-cohesin interactions and chromatin colocalization are not disrupted in ΔZF1 mutants, related to. Figure 3. (A, B) Volcano plot of CTCF differential protein interactions comparing ESC WT vs (A) ESC-ΔZF1 or (B) ESC-ΔZF10 (see all in **Table S1** or **Table S2**, respectively). Flag-Halo-tagged CTCF was purified by anti-Flag immunoprecipitation in native conditions and protein interactions were identified by MS. Adjusted p-value cutoff: ≤ 0.05, log_2_ fold-change cutoff: ≥ 1, ≤-1, N=2. (C, D) DiffBind MA plot of differentially called (C) Rad21 and (D) Smc3 ChIP-seq peaks comparing ESC- ΔZF1 vs WT. Adjusted p-value cutoff: ≤ 0.05, log_2_ fold-change cutoff: ≥ 1, ≤-1, N=2. (E, F) DiffBind MA plot of differentially called (E) Rad21 and (F) Smc3 ChIP-seq peaks comparing NPC- ΔZF1 vs WT. Adjusted p-value cutoff: ≤ 0.05, log_2_ fold-change cutoff: ≥ 1, ≤-1, N=2. (G, H) ChIP-seq heatmaps of (G) Rad21 and (H) Smc3, comparing chromatin binding in ESC-ΔZF1 vs WT. Each row is a loop anchor coordinate, and the heatmap is clustered based on whether the anchors are ΔZF1-lost or ΔZF1-retained. (I, J) ChIP-seq heatmaps of (I) Rad21 and (J) Smc3, comparing chromatin binding in NPC-ΔZF1 vs WT. Each row is a loop anchor coordinate, and the heatmap is clustered based on whether the anchors are ΔZF1-lost or ΔZF1-retained.

**Figure S7.**
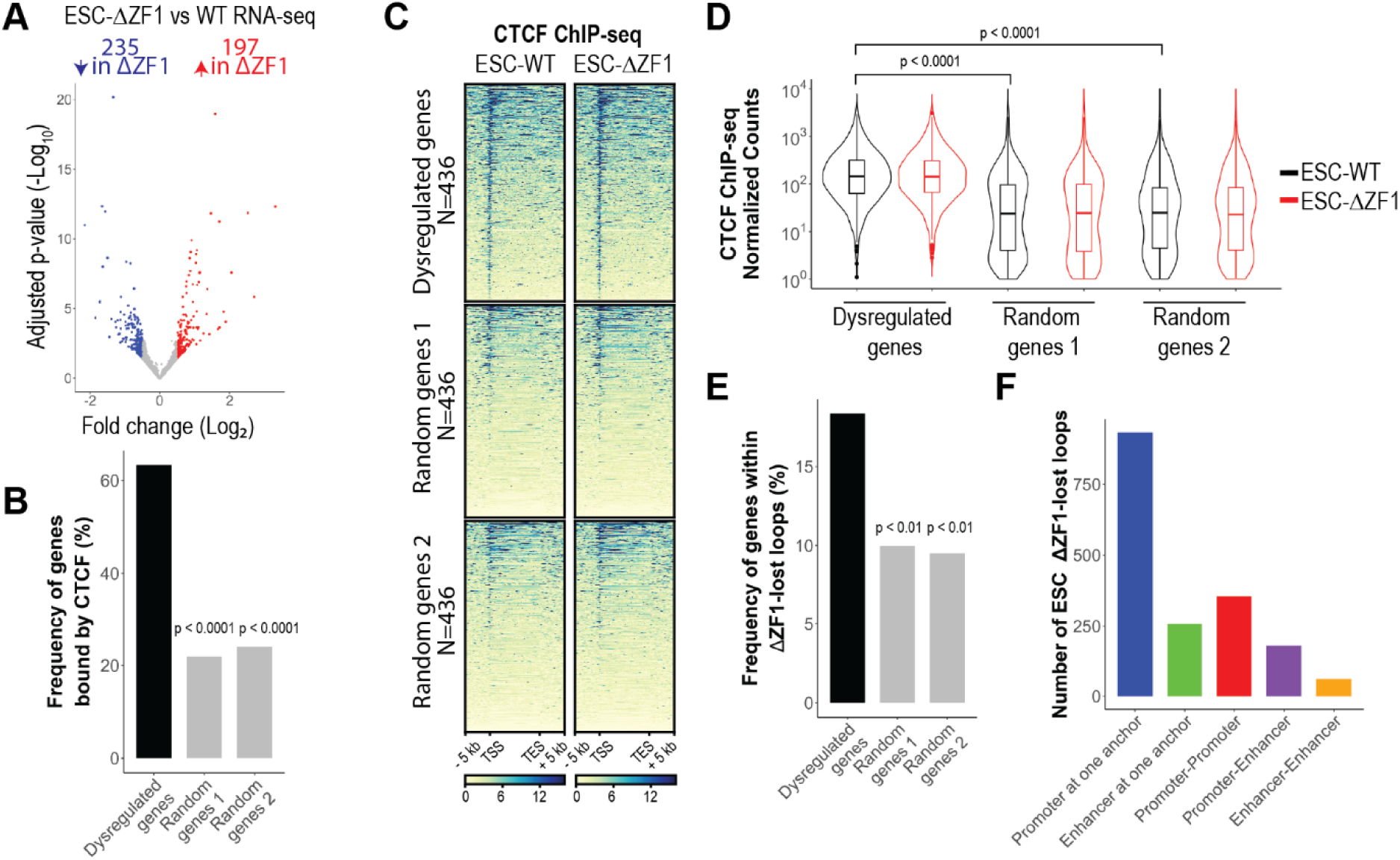
Dysregulated genes in ESC-ΔZF1 mutant are enriched at disrupted loops, related to. Figure 4. (A) Deseq2 volcano plots showing gene expression changes in ESC-ΔZF1 compared to ESC-WT (see all in **Table S3**). Adjusted p-value cutoff: ≤ 0.05, log_2_ fold-change cutoff: ≥ 0.5, ≤-0.5, N=2. (B) Bar plots showing the percentage of genes from each gene set (dysregulated, random genes 1, random genes 2) that overlap with CTCF ChIP-seq peaks in ESCs. p-values comparing each of the random gene sets to dysregulated genes were determined using Fisher’s Exact test. (C) CTCF ChIP-seq heatmaps at the TSS to TES of dysregulated genes and two sets of randomly generated genes in ESCs. (D) Violin plots quantifying CTCF ChIP-seq reads from TSS to TES of genes shown in (C). p-values were determined using t-test. (E) Bar plots showing the percentage of genes from each gene set (dysregulated, random genes 1, random genes 2) that are co-localized within ΔZF1-lost loops in ESCs. p-values comparing each of the random gene sets to dysregulated genes were determined using Fisher’s Exact test. (F) Bar plots showing the number of ESC-ΔZF1-lost anchors that overlapped with promoters and/or enhancers.

**Figure S8.**
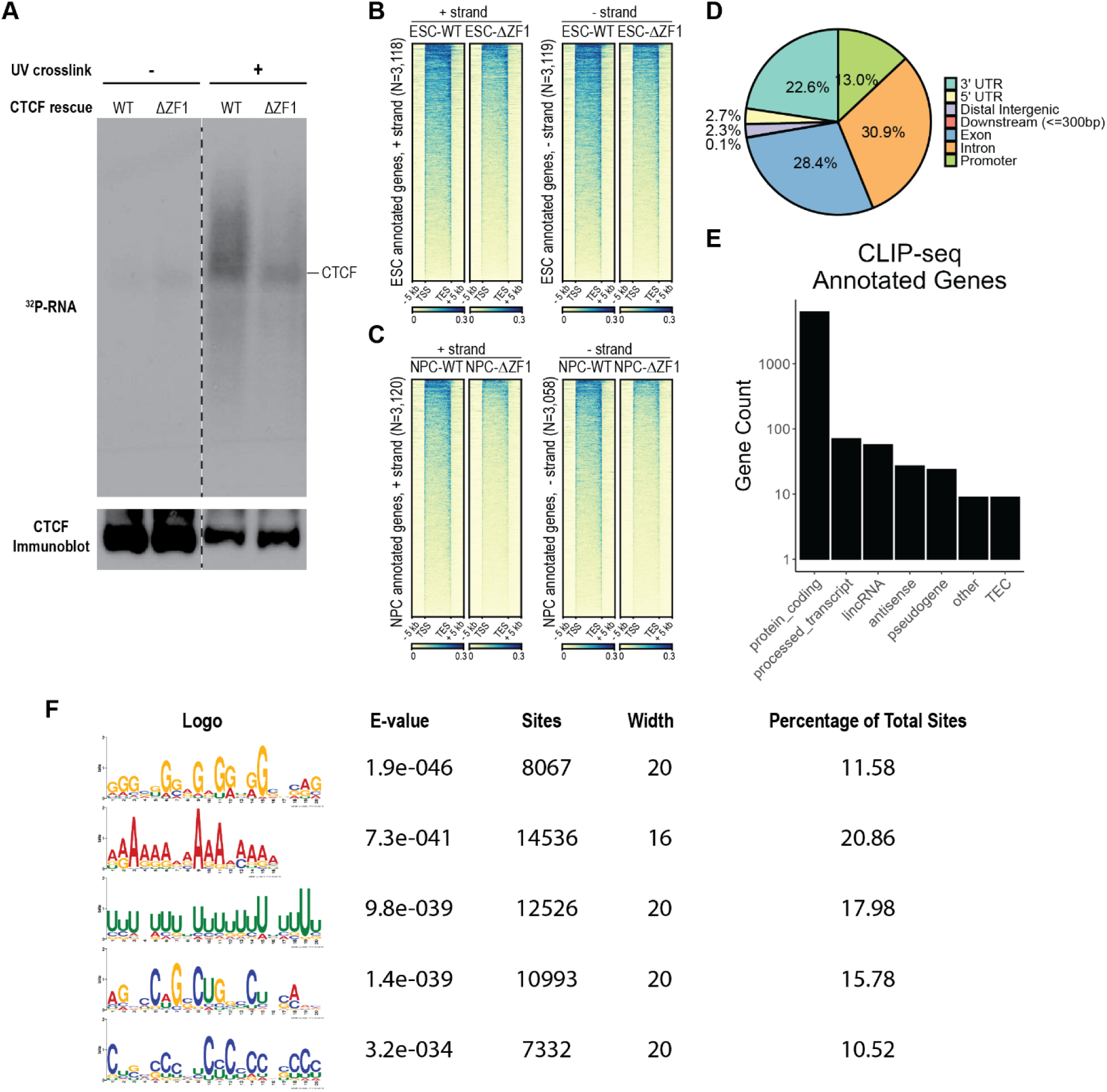
CTCF CLIP-seq gene annotations and characteristics, related to. Figure 5. (A) Isotope labeling of CTCF RNA interactome and immunoblot of CTCF. Flag-halo-tagged WT and ΔZF1 CTCF were immunoprecipitated with anti-flag beads under non-crosslinking and UV-crosslinking conditions. RNAs purified from CTCF immunoprecipitation were labeled using radioactive ATP. The labeled protein-RNA complexes were separated by gel electrophoresis and transferred onto nitrocellulose membrane. Membrane was exposed to film to visualize ^32^P-labeled RNA. CTCF western blot was done on the same membrane. Dashed line indicates exclusion of unused lanes on the membrane. (B, C) WT and ΔZF1 CLIP-seq heatmaps at CLIP-seq annotated genes in (B) ESCs and (C) NPCs. (D) ESC-WT CLIP peaks were annotated to genomic features using ChIPseeker. Pie chart shows the percentages of CLIP peaks annotated to specific features. (E) ESC-WT CLIP peaks were annotated to genes and gene types. Bar chart shows the number of CLIP annotated genes for different gene types. (F) RNA Motif identification for ESC-WT CLIP peaks using MEME suite. RNA-seq reads were used as background sequences. Motif search in MEME was performed *de novo* until 5 motifs were found.

**Figure S9.**
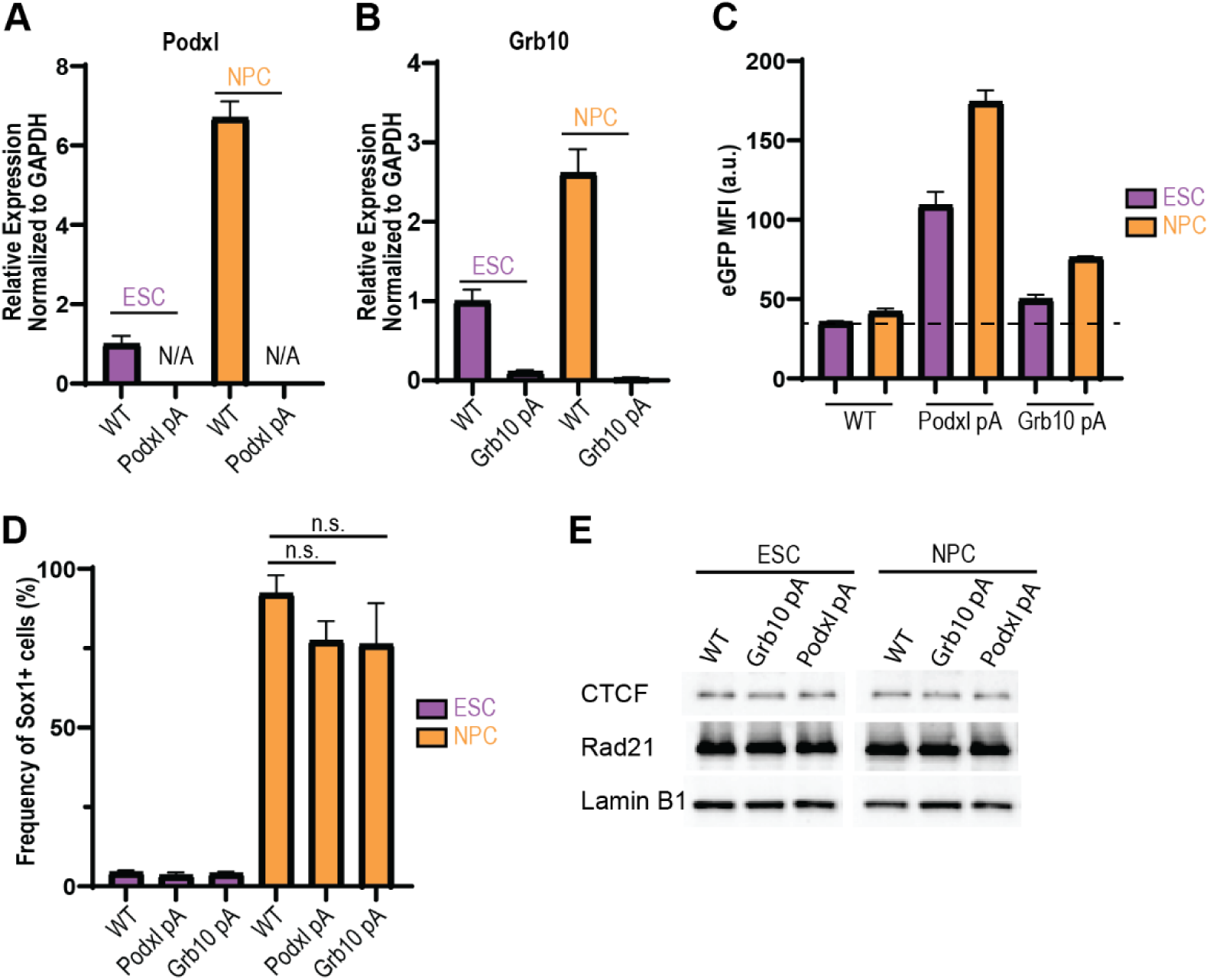
Functional evaluation of Podxl-T2A-eGFP-SV40pA and Grb10-T2A-eGFP-SV40pA RNA- truncation mutants, related to. Figure 7. (A-B) WT and (A) Podxl pA and (B) Grb10 pA cells were maintained as ESCs or differentiated into NPCs. RT-qPCR was done on (A) *Podxl* gene and (B) *Grb10* gene downstream of the pA insertion and normalized with GAPDH. Data are represented as mean ± SEM, N=3, N/A: Cq values > 40. (C) eGFP fluorescence of the Podxl pA and Grb10 pA mutants were analyzed by flow cytometry. Bar plots show mean fluorescence intensity (MFI, arbitrary units) of eGFP in the parental WT, Podxl pA, and Grb10 pA clones. Data are represented as mean ± SEM, N=2-3. Dashed line is the baseline fluorescence for eGFP-negative WT cells. (D) NPC differentiation was assessed in the pA mutants in comparison to the parental WT cells. Cells were fixed and immunostained with anti-Sox1 antibody. The percentages of Sox1+ cells were then analyzed by flow cytometry. Data are represented as mean ± SEM, p-values were determined using Dunnett’s multiple comparison test, n.s.=not significant, N=2-3. (E) Western blot of CTCF and Rad21 in WT, Grb10 pA, and Podxl pA cells.

